# The Greatwall/PP2A-B55α axis remodels the G2/M boundary and shapes cellular dependence on PKMYT1

**DOI:** 10.64898/2026.09.04.749407

**Authors:** Róbert Zach, Alex D. Herbert, Calin-Mihai Dragoi, Megan Meredith, Lisandra Gadelkarim, William R. Foster, Helfrid Hochegger

## Abstract

The switch-like G2/M transition and mitotic exit depend on a feedback loop comprising CDK1, Greatwall kinase, and the PP2A-B55α phosphatase. Monogenic disruptions of these regulators impair cellular functions, drive genomic instability, and promote cancer-associated characteristics. Yet how perturbations of this feedback loop interact, and whether their effects depend on cellular context, remain unclear. Here, we assess how combinatorial perturbations of CDK1, Greatwall, and PP2A-B55α affect cell-cycle control in non-transformed RPE-1 and tumour-derived HeLa cells. We identify PP2A-B55α as a context-dependent G2/M repressor with a critical role in cancer cells, where it cooperates with PKMYT1 and, to a lesser extent, WEE1 to prevent premature mitosis. In cancer cells, but not in non-transformed cells, reduced PP2A-B55α activity and Greatwall overexpression produce marked sensitisation to the PKMYT1-selective RP-6306 and modest sensitisation to the WEE1-selective adavosertib. This asymmetric sensitisation reflects a functional separation between WEE1 and PKMYT1 in S- and G2-phase control. Our work outlines fundamental and cell-type-specific contributions of CDK1, Greatwall and PP2A-B55α to cell-cycle regulation and proposes G2/M plasticity as a vulnerability with utility in PKMYT1-targeted therapy.

## Introduction

Transitions between cell-cycle phases are orchestrated by bistable feedback mechanisms that regulate the switch between inactive and active states of cyclin-dependent kinases (CDKs) ^1,2^. A classic example of this principle is mitotic entry, which is driven by a rapid rise in CDK1 activity ^3,4^. Throughout interphase, CDK1-dependent phosphorylation is restrained by two complementary mechanisms. First, WEE1 and PKMYT1 catalyse inhibitory phosphorylation of CDK1 at tyrosine 15 (Y15) and threonine 14 (T14), respectively ^5–8^. Second, PP2A-B55 phosphatases dephosphorylate CDK1 substrates ^9–11^. At the point of mitotic entry, CDC25 phosphatases activate CDK1 by removing inhibitory phosphates from T14 and Y15 ^12–14^. CDK1 then reinforces its own activation through two positive feedback loops. First, CDK1 directly phosphorylates and inactivates its inhibitors, WEE1 and PKMYT1 ^15,16^. Second, CDK1 activates the Greatwall (GWL) kinase ^17,18^, which phosphorylates ENSA and ARPP19 at serine 67 (S67) and serine 62 (S62), respectively ^19,20^. Phosphorylated ENSA(S67) and ARPP19(S62) then inhibit PP2A-B55 phosphatases with high affinity and inhibit their activity through an unfair competition mechanism ^21,22^. These two positive feedback loops establish a bistable switch, ensuring switch-like mitotic commitment. Their activation favours the transition to a state of high CDK1 and low PP2A-B55 activity and thus, the stable phosphorylation of CDK1 substrates required for faithful cell division.

In mitosis, CDK1 phosphorylates more than 1,000 proteins that drive nuclear envelope breakdown, chromosome condensation, and mitotic spindle assembly ^23–25^. Notably, the stability of the mitotic phosphoproteome depends on continuous phosphorylation of ENSA(S67) and ARPP19(S62), which sequester PP2A-B55 phosphatases and limit their activity towards CDK1 substrates ^19–22,26^. During mitotic exit, the Anaphase-Promoting Complex/Cyclosome (APC/C) promotes degradation of Cyclin B, the essential CDK1 cofactor, leading to a decline in CDK1 activity, inactivation of GWL, and dephosphorylation of ENSA(S67) and ARPP19(S62) ^27–29^. As a result, PP2A-B55 complexes, no longer occupied by their pseudo-substrates, drive the orderly reversal of CDK1-dependent phosphorylation events, ensuring that chromosome segregation occurs before nuclear envelope reformation and cytokinesis ^10,30,31^.

Perturbations in the CDK1/GWL/PP2A-B55 axis have primarily been studied by targeting its individual components. Depending on the extent, inhibition of CDK1 has been associated with defects in mitotic outcomes or with failure to enter mitosis, which, in some contexts, can lead to unscheduled APC/C activation and endoreduplication ^32–38^. Genetic or pharmacological disruption of GWL destabilises mitotic phosphorylation, resulting in mitotic catastrophe or defective cytokinesis ^26,39–42^. Finally, loss of the regulatory PP2A subunit B55 disrupts the ordered dephosphorylation programme during mitotic exit, impairing the chronology of chromosome segregation, nuclear envelope reformation, and cytokinesis ^9,10,43^. The B55 family comprises four paralogues (B55α-δ) that regulate overlapping but distinct substrate pools in a context-dependent manner ^44^. For instance, in mice, B55α plays an essential role in embryonic development, but its dysfunction can be compensated for by B55δ in mouse embryonic fibroblasts ^43^. Importantly, in HeLa and RPE-1 human cell line models, depletion of B55α alone fully suppresses the phenotypes caused by GWL depletion or inhibition, suggesting that, in these contexts, PP2A-B55α is the predominant contributor to the GWL signalling cascade ^26^.

Many tumour tissues overexpress CDK1 and its positive regulators, which are thought to weaken DNA damage checkpoints and promote uncontrolled proliferation and aggressive tumour growth ^45–48^. Elevated GWL expression has also been reported across a broad range of cancers, contributing to chromosomal instability, cellular transformation, partial epithelial-mesenchymal transition, metastatic relapse, and poor prognosis ^49–52^. Finally, loss of the tumour-suppressing B55α has been linked to defective homologous recombination, micronucleus formation, and reduced survival in patients with breast, prostate, and non-small cell lung cancers ^53–59^.

Although perturbation of individual components of the CDK1/GWL/PP2A-B55 axis has been extensively studied, the consequences of distinct combined dysregulations across multiple nodes of the pathway remain largely unexplored. Additionally, the frequent deregulation of this signalling axis in human cancers raises questions about the compensatory mechanisms that enable tumour cells to tolerate such perturbations and whether these adaptations expose therapeutic vulnerabilities. To address this, we established an experimental system that enables simultaneous perturbation of CDK1, GWL, and PP2A-B55α and allows dissection of their individual contributions and interactive effects on cell-cycle control and mitotic fidelity in both non-transformed and cancer-derived cell line models.

## Results

### Tumours dysregulate the balance of CDK1, GWL, and B55α expression

To assess the extent of tumour-specific expression-level imbalances within the CDK1/GWL/PP2A-B55α axis, we analysed RNA expression levels (FPKM) of *CDK1*, *MASTL* (*GWL*), and *PPP2R2A* (*B55α*) across tumour-adjacent normal tissues and tumour-derived resections using The Cancer Genome Atlas (TCGA) data. For each gene pair (^Y^*GWL vs* ^X^*CDK1*, ^Y^*B55α vs* ^X^*CDK1*, and ^Y^*B55α vs* ^X^*GWL*), we quantified individual gene expression levels, pairwise expression relationships, and expression-level imbalances, expressed as relative dominance scores (X - Y) / (X + Y), ranging from -1 (complete dominance of gene Y) to +1 (complete dominance of gene X) (Fig. 1a-c). Tumours showed markedly dysregulated expression of all three genes, with pairwise co-expression signatures extending beyond the ranges observed in normal tissues (Fig. 1a-d). This spread was particularly evident in *CDK1*-containing pairs, where *CDK1* levels frequently exceeded those of *GWL* and *B55α,* shifting relative dominance scores towards +1 (Fig. 1, a, b, and e). The combined distribution of *GWL* and *B55α* expression levels showed less pronounced broadening, with a moderate skew towards relative dominance scores > 0, indicative of disproportionately elevated *GWL* relative to *B55α* expression (n = 1,354 tumour biopsies, 13.2%) (Fig. 1c-e and Supplementary Fig. S1). While *CDK1*-dominant co-expression cases were also present in a subset of normal tissues, the relative dominance of *GWL* over *B55α* was observed only in tumour samples, highlighting it as a putative cancer-specific signature (Fig. 1e and Supplementary Fig. S1). Although transcript levels are imperfect predictors of stoichiometry and pathway functionality, these results indicated marked variation in the relative abundance of CDK1/GWL/PP2A-B55α components across tumour tissues, with likely implications for their functional interactions and regulatory roles within broader signalling contexts.

**Fig. 1.**
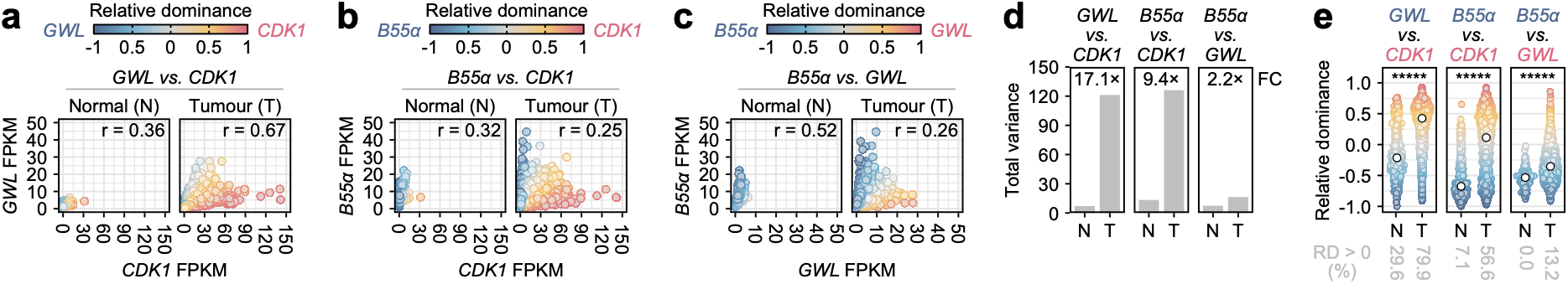
Pan-cancer TCGA RNA expression profiling of *CDK1*, *GWL* and *B55α*. **(a-c)** Scatterplots showing RNA expression levels (FPKM) of *CDK1*, *MASTL* (*GWL*) and *PPP2R2A* (*B55α*), and their pairwise relationships, *GWL vs CDK1* (a), *B55α vs CDK1* (b) and *B55α vs GWL* (c), in pan-cancer tumour-adjacent normal (N; 23 tissue types, n = 723) and tumour (T; 33 cancer types; n = 10,296) samples. Colour gradients represent relative dominance (RD) values, calculated as RD = (x - y) / (x + y), where x and y are the gene expression values on the x- and y-axes, respectively. Positive values (warm colours) indicate relative dominance of the gene on the x-axis; negative values (cool colours) indicate relative dominance of the gene on the y-axis. r – Pearson correlation coefficient. **(d)** Bar plots showing the trace of the covariance matrix (total variance) for the three analysed gene pairs, quantifying the overall spread of pairwise expression states, calculated as: total variance = Var(x) + Var(y), where Var(x) and Var(y) represent the variances of the expressions on the x- and y-axes, respectively. Fold change (FC) was calculated as FC = T / N, where T and N represent the total variances of tumour and normal samples, respectively. **(e)** Distributions of RD scores shown as density-scaled jittered points. Colour-coded points and white points represent the relative dominance values of individual samples and their medians, respectively. Statistical significance was assessed using a two-sided Wilcoxon rank-sum test, with reported p-values uncorrected. ***** p < 0.00001.

### Cell-cycle-focused analysis of CDK1, GWL, and PP2A-B55α perturbations

Although cancer cells frequently dysregulate the CDK1/GWL/PP2A-B55α axis, how combined perturbations of its components affect cell-cycle control remains poorly understood. To address this, we developed a protocol to simultaneously perturb CDK1, GWL, and PP2A-B55α and assess the resulting effects on interphase progression and mitotic fidelity in tumour-derived HeLa (cervical carcinoma) and non-transformed, hTERT-immortalised RPE-1 cells. We modulated CDK1 activity with two sub-saturating doses (2 µM and 4 µM) of the CDK1 inhibitor RO-3306 ^32^, which reduced prometaphase phosphorylation of PRC1(T481) to 86% and 62% in HeLa cells, and to 29% and 11% in RPE-1 cells, respectively (Fig. 2a, b and Supplementary Fig. S2a-f). Although these changes indicated varying degrees of CDK1 inhibition between the two cell lines, because PRC1(T481) phosphorylation reflects opposing activities of CDK1 and PP2A-B55α, we could not reliably infer the extent of CDK1 inhibition per se. We inhibited GWL using a saturating dose (2 µM) of the selective GWL inhibitor C-604 ^26^, which reduced the relative phosphorylation of the canonical GWL substrates ENSA(S67) and ARPP19(S62) to < 10% of control levels in both cell lines (Fig. 2a-c and Supplementary Fig. S2g-j). Finally, we used a pool of B55α-specific siRNAs (siB55α), which reduced B55α expression to approximately 25% of endogenous levels 48 h after siRNA transfection (Fig. 2a, d and Supplementary Fig. S2k).

**Fig. 2.**
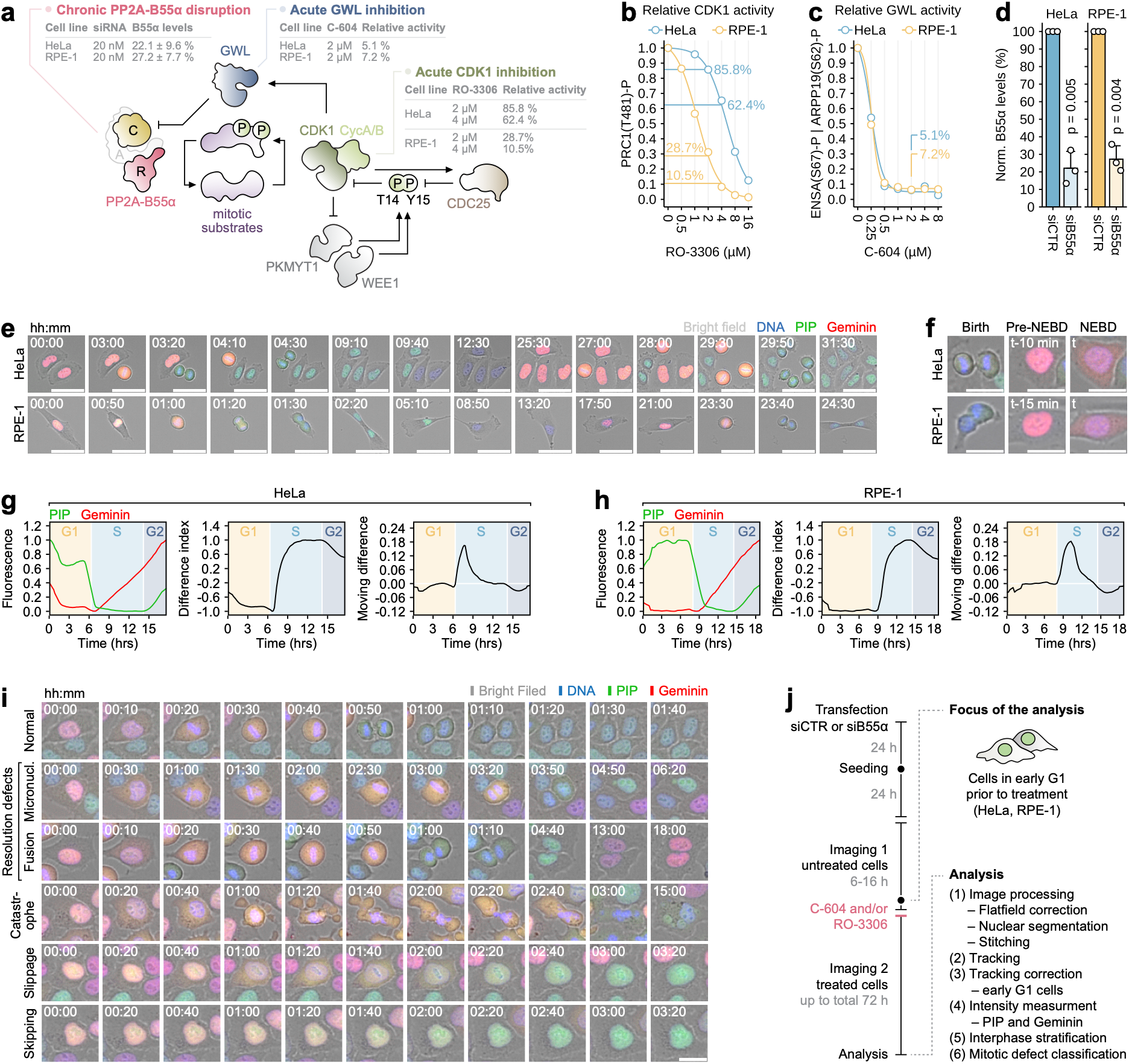
An experimental system to study multifactorial perturbation of CDK1, GWL and PP2A-B55α. **(a)** Simplified schematic of the CDK1/GWL/PP2A-B55α signalling circuit, including auxiliary regulators of CDK1 kinase activity, WEE1 and PKMYT1. Experimental perturbations of CDK1, PP2A-B55α and GWL (B-D) are indicated. **(b)** The impact of RO-3306 on relative CDK1 activity in HeLa and RPE-1 cells, inferred from dose-dependent changes in PRC1(T481) phosphorylation. Points and lines represent the means of n = 3 biological replicates and fitted four-parameter sigmoidal models, respectively. Inhibition levels at 2 µM and 4 µM RO-3306 are shown. **(c)** The effect of C-604 on relative GWL activity in HeLa and RPE-1 cells, inferred from changes in ENSA(S67) and ARPP19(S62) phosphorylation. Points and lines represent the means of n = 2 biological replicates and fitted four-parameter models, respectively. The inhibition level at 2 µM C-604 is shown. **(d)** B55α protein levels quantified by western blot densitometry (relative to histone H3) in HeLa and RPE-1 cells 48 hours after transfection with scrambled control (siCTR) and B55α-targeting (siB55α) siRNA. Bars and error bars represent means and standard deviations, respectively. Points represent results from n = 3 biological replicates. Statistical significance was assessed using an unpaired two-tailed t-test, with p-values indicated. **(e)** Representative time-lapse images of live HeLa and RPE-1 PIP-FUCCI cells. Scale bars represent 50 µm. **(f)** Representative images of cell birth and mitotic entry, marked by nuclear envelope breakdown (NEBD), in HeLa and RPE-1 PIP-FUCCI cells. Scale bar is 25 µm. **(g-h)** Representative PIP-FUCCI track analyses in HeLa (g) and RPE-1 (h) cells. Left panels – normalised mean nuclear fluorescence intensities of PIP-eGFP and Geminin-mCherry probes. Middle panels – normalised difference index (DI), calculated as DI = (Geminin – PIP) / (Geminin + PIP). Right panels – moving difference (slope) using a 9-frame window of the normalised and scaled DI values. **(i)** Representative images of normal and pathological mitotic outcomes, including resolution defects, catastrophes, slippages and skipping events, in HeLa PIP-FUCCI cells. Scale bar is 25 µm. **(j)** Schematic representation of the complete experimental procedure.

To assess how these perturbations affected cell-cycle progression, we employed the live-cell PIP-FUCCI system, comprising two cell-cycle probes, PIP-eGFP and Geminin-mCherry ^60,61^ (Fig. 2e). Phase-specific expression of these probes, together with single-cell tracking and computational image analysis, enabled us to partition each interphase (from cell birth to mitotic entry) into G1, S, and G2 phases and to quantify their durations (Fig. 2f-h and Supplementary movie M1). Additionally, PIP-FUCCI live-cell imaging enabled classification of distinct mitotic outcomes, including mitotic resolution defects (such as micronuclei, daughter-cell fusions, persistent cytoplasmic bridges, and post-mitotic cell-cycle arrests), as well as mitotic catastrophe, mitotic slippage, and mitotic skipping (Fig. 2i, Supplementary Figs. S3 and S4 and Supplementary movies M2 and M3).

Each experiment comprised two consecutive live-cell imaging segments (Fig. 2j). In the first segment (up to 16 h), we imaged siCTR- or siB55α-transfected HeLa and RPE-1 PIP-FUCCI cells to determine the cell-cycle phase of each cell in the field of view (Fig. 2j). We then removed the cells from the microscope and treated them with RO-3306, C-604, or their combination. The second imaging segment (up to 72 h) began immediately after treatment and captured acute cellular responses to six combinations of RO-3306 (0, 2, or 4 µM) and C-604 (0 or 2 µM) under conditions of normal or reduced B55α expression (Fig. 2j). Notably, splitting the experiment into two imaging segments allowed us to analyse cells in early G1 at the time of treatment, which was necessary to characterise the effects of CDK1/GWL/PP2A-B55α perturbations on the entire interphase and mitosis.

### Perturbations of CDK1, GWL and PP2A-B55α produce conserved and cell-type-specific interphase phenotypes

Acute CDK1 inhibition alone did not alter G1 length in either cell line, confirming the established view that CDK1 is not rate-limiting for G1 progression (Fig. 3a and Supplementary Fig. S5). Conversely, in line with prior reports ^9^, reduced B55α expression increased G1 duration in both HeLa (mean ± SD; siCTR: 417 ± 12 min, siB55α: 573 ± 33 min) and RPE-1 (siCTR: 338 ± 8 min, siB55α: 443 ± 34 min) cells (Fig. 3a and Supplementary Fig. S5). This delay was suppressed by acute CDK1 inhibition in RPE-1 but not in HeLa cells, suggesting that PP2A-B55α could promote G1 progression through multiple cell-type-specific mechanisms (Fig. 3a and Supplementary Fig. S5). Acute GWL inhibition with 2 µM C-604 did not alter G1 duration in HeLa cells but produced a marked G1 delay in RPE-1 cells that was not reversed by PP2A-B55α depletion (siCTR: 535 ± 48 min, siB55α: 645 ± 141 min). Surprisingly, co-treatment with RO-3306 also attenuated this delay, implying a yet-uncharacterised cell-type-specific role of GWL/CDK1 signalling during G1 (Fig. 3a and Supplementary Fig. S5). S-phase was affected only by 4 µM RO-3306, which prolonged it in both HeLa (mean ± SD; 0 µM: 490 ± 30 min, 4 µM: 620 ± 35 min) and RPE-1 (0 µM: 400 ± 43 min, 4 µM: 470 ± 30 min) cells (Fig. 3b and Supplementary Fig. S5). This effect was independent of GWL or PP2A-B55α activity and likely reflected either the role of CDK1 in DNA synthesis ^34,62^ or off-target inhibition of CDK2 by RO-3306 ^32^. Overall, these results confirmed that CDK1 activity is not rate-limiting for G1 progression and suggested that, at least in HeLa and RPE-1 cells, GWL and PP2A-B55α do not contribute to S-phase control. Additionally, this analysis revealed putative cell-type-specific contributions of PP2A-B55α and GWL to G1 regulation, warranting further experimental validation.

**Fig. 3.**
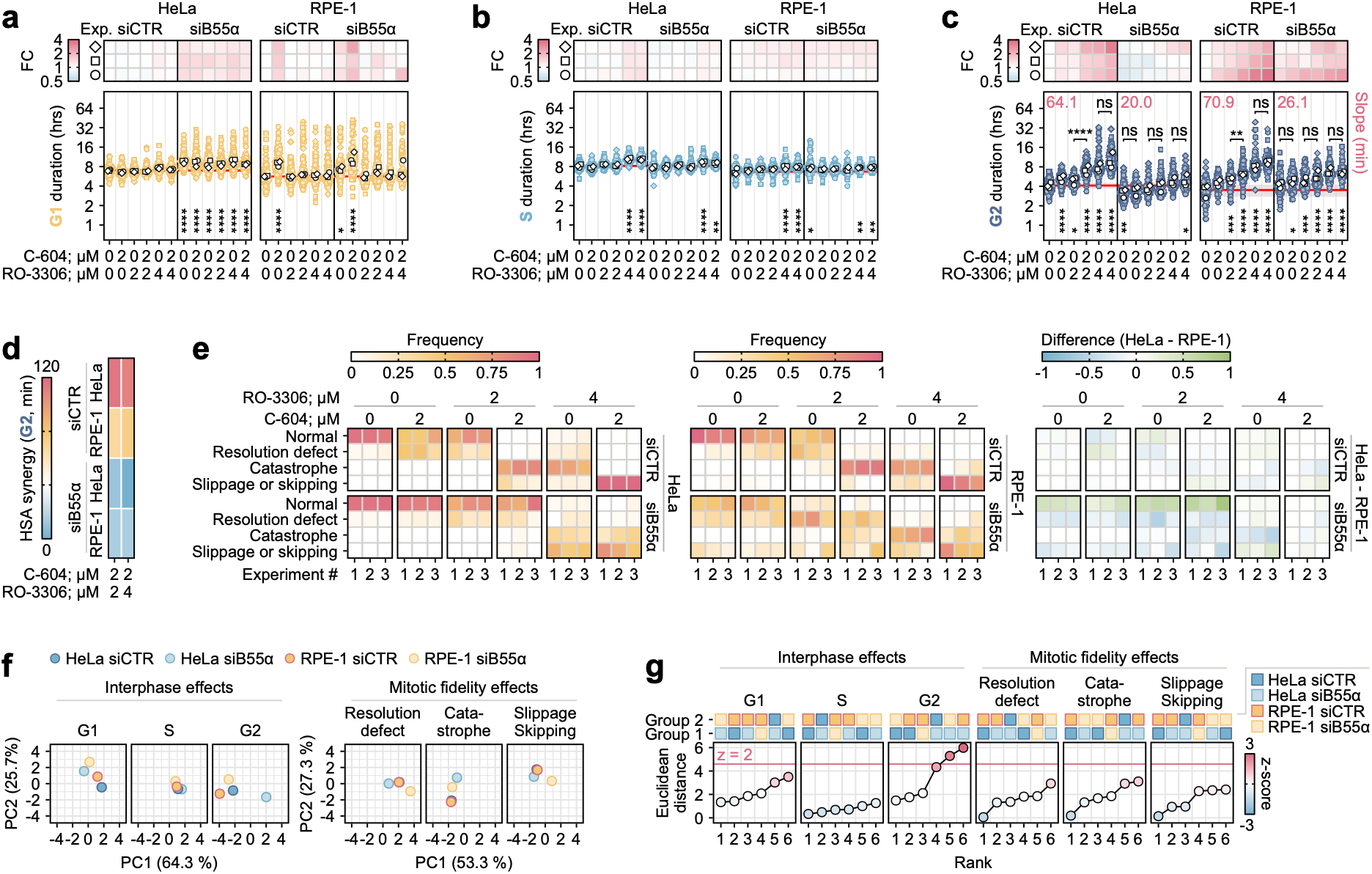
Cell-cycle effects of single and combined perturbations of CDK1, GWL and PP2A-B55α. **(a-c)** Quantification of interphase effects in HeLa and RPE-1 PIP-FUCCI cells transfected with control (siCTR) or B55α-targeting (siB55α) siRNAs and treated with C-604 (2 µM) and/or RO-3306 (2 or 4 µM) as indicated. Bottom panels – Quantification of G1 (a), S (b) and G2 (c) durations. Coloured and white points represent individual measurements and the medians of n = 3 biological replicates, respectively. Horizontal red lines and accompanying shaded regions represent means and standard deviations computed from three independent medians of siCTR-transfected cells treated with DMSO (reference). Top panels – Heatmaps displaying the median fold changes (FC) in G1, S and G2 lengths relative to the reference. Symbols (circle, square, diamond) indicate results of individual biological replicates. Numerical values above each plot indicate the slope of the linear regression fitted across conditions (ordered by increasing perturbation strength), quantifying the overall steepness of phase-duration responses. For each biological replicate, between 10 and 52 mitotic events were scored for each group. Statistical significance was assessed using the Kruskal-Wallis test, followed by Dunn’s post hoc test with a Bonferroni correction. A more stringent statistical significance threshold was used: * p < 0.01, ** p < 0.001, *** p < 0.0001, **** p < 0.00001. **(d)** A heatmap showing the Highest Single Agent (HSA) synergy scores between the effects of RO-3306 (2 or 4 µM) and C-604 (2 µM) on G2 phase duration, calculated from overall medians. HSA scores > 0 indicate synergy; ∼ 0 indicates no additional effect beyond the strongest single agent; < 0 indicates antagonism. **(e)** Left and middle panels – Heatmaps showing the frequencies (0-1) of distinct classes of mitotic outcomes in HeLa (left) and RPE-1 (middle) cells transfected with control (siCTR) or B55α-targeting (siB55α) siRNAs and treated with C-604 (2 µM) and/or RO-3306 (2 or 4 µM). Mitotic outcomes were scored manually and classified as: normal, resolution defect (micronuclei, daughter cell fusions, cytoplasmic bridge, post-mitotic cell-cycle exit), catastrophe, and a joint category of slippage or skipping. Results from n = 3 biological replicates are shown. Between 10 and 52 mitotic events were scored per condition per biological replicate, with variation in counts primarily reflecting cell-tracking limitations rather than treatment effects. Right panel – Heatmaps showing differences in the occurrence of distinct mitotic outcomes between HeLa and RPE-1 cells (HeLa minus RPE-1; range -1 to 1). Positive values indicate outcomes are more frequent in HeLa; negative values indicate outcomes are more frequent in RPE-1; zero indicates equal frequencies. **(f)** Principal component analysis (PCA) projections of treatment responses (RO-3306 at 0, 2, 4 µM × C-604 at 0, 2 µM) in four groups (HeLa siCTR, HeLa siB55α, RPE-1 siCTR, RPE-1 siB55α) onto a two-dimensional space, performed separately for interphase phenotypes (G1, S, G2; mean fold-change relative to untreated control) and for frequencies of mitotic outcomes (resolution defects, catastrophes, and slippage/skipping events, with normal events excluded). Each point represents a (group × phenotype) combination, characterised by its six-dimensional response vector across drug treatments. PC1 and PC2 are shown, with the percentage of variance explained indicated on each axis. **(g)** Ranked pairwise Euclidean distances between groups for each phenotypic output, computed in PC1-PC2 space. Z-scores for 36 distances (six group-wise comparisons for six phenotypes) were computed across the combined distribution. Higher z-scores indicate greater divergence between groups relative to the overall distribution of pairwise distances.

Consistent with the established roles of CDK1 and GWL in promoting the G2/M transition, acute single-agent RO-3306 or C-604 treatment increased G2 duration in both HeLa and RPE-1 cells (Fig. 3c and Supplementary Fig. S5). In both cell lines, the combination of RO-3306 and C-604 prolonged G2 beyond the effect of either inhibitor alone, as reflected by positive highest single-agent (HSA) synergy scores (Fig. 3c, d; fig. S5). Notably, reduced PP2A-B55α activity had two qualitatively distinct effects. (1) In both HeLa and RPE-1 cells, B55α depletion attenuated the G2-prolonging effects of single-agent RO-3306 and C-604 treatments, as well as their synergy, resulting in flattened dose-response profiles across the full treatment matrix (Fig. 3c, d and Supplementary Fig. S5). PP2A-B55α therefore acted as a conserved driver of G2 delay in cells with reduced CDK1 and GWL activity. (2) B55α depletion significantly reduced baseline G2 duration in HeLa cells (siCTR: 247 ± 26 min; siB55α: 192 ± 28 min) but not in RPE-1 cells (siCTR: 210 ± 46 min; siB55α: 255 ± 27 min) (Fig. 3c and Supplementary Fig. S5). Thus, in unperturbed cells, PP2A-B55α contributed to G2/M regulation in a cell-type-specific manner. Collectively, these results broadly aligned with the bistability model and confirmed that the G2/M switch serves as a point of convergence, in which GWL promotes full CDK1 activation by inhibiting PP2A-B55α. Interestingly, we also revealed that, under normal conditions, PP2A-B55α contributed to the timely G2/M transition only in HeLa cells, implying cell-type-specific differences in how PP2A-B55α integrates into the wider G2/M control network.

### Perturbations of CDK1, GWL, and PP2A-B55α produce discrete mitotic phenotypes

Although mitotic commitment is a switch-like event triggered when CDK1 activity exceeds a critical threshold, some evidence indicates that post-commitment CDK1 activity continues to rise before declining at the onset of anaphase ^4,63^. This progressive activation is thought to establish discrete CDK1 activity thresholds that coordinate the temporal ordering and execution of mitotic events. Consistent with this quantitative model of mitotic progression, single and combined perturbations of the CDK1/GWL/PP2A-B55α axis produced discrete mitotic phenotypes, indicative of stage-specific points of mitotic failure (Supplementary movies M2, M3).

HeLa and RPE-1 cells with reduced CDK1 activity (siCTR; 2 µM RO-3306, 0 µM C-604) or abolished GWL activity (siCTR; 0 µM RO-3306, 2 µM C-604) exhibited a higher incidence of mitotic resolution defects, including micronuclei, incomplete separation of daughter cells, and, in RPE-1 cells, post-mitotic cell-cycle arrest (Fig. 3e). These findings indicated that reduced CDK1 activity or near-complete GWL inhibition allowed progression through early mitosis but compromised later events required for the resolution of daughter cells and their nuclei. Consistent with stage-specific requirements for mitotic CDK1 activity, further CDK1 inhibition (siCTR; 4 µM RO-3306, 0 µM C-604) shifted the point of mitotic failure to earlier, pre-anaphase stages. This defective exit manifested as mitotic catastrophe, characterised by an inability to segregate mitotic chromosomes, defective cytokinesis, and nuclear fragmentation (Fig. 3e and Supplementary Fig. S4). A combination of partial CDK1 inhibition and abolished GWL activity (siCTR; 2 µM RO-3306, 2 µM C-604) produced mitotic catastrophe comparable to that induced by 4 µM RO-3306 alone, demonstrating that GWL inhibition synergistically reduced effective CDK1 activity, most likely through derepression of the CDK1-antagonistic PP2A-B55α (Fig. 3e). In further support of graded, stage-specific requirements for mitotic CDK1 activity, simultaneous inhibition of CDK1 and GWL (siCTR; 4 µM RO-3306, 2 µM C-604) typically resulted in mitotic slippage or skipping, characterised by untimely APC/C activation (marked by Geminin degradation) and early reversal of the G2/M state, ultimately resulting in a single G1 cell with an intact nucleus and duplicated DNA content (Fig. 3e and Supplementary Fig. S4). Collectively, these findings indicated that a progressive reduction in effective CDK1 activity, whether achieved directly or via GWL inhibition and PP2A-B55α derepression, produced conserved, stage-specific mitotic pathologies, consistent with a proposed model in which discrete thresholds of CDK1 activity govern successive steps of mitotic progression ^4,63^.

While B55α depletion alone did not disrupt mitotic progression in HeLa cells, it increased the incidence of mitotic resolution defects, mitotic slippage, and skipping in RPE-1 cells (siB55α; 0 µM RO-3306, 0 µM C-604) (Fig. 3e). This suggested that successful mitotic completion was more dependent on PP2A-B55α activity in RPE-1 than in HeLa cells. Consistent with PP2A-B55α derepression as a key mediator of the cytotoxic effects of GWL inhibition ^26,64^, B55α depletion mitigated the deleterious effects of C-604 (siB55α; 0 µM RO-3306, 2 µM C-604) and reduced the synergistic interaction between C-604 and RO-3306 in both cell lines (siB55α; 2 µM RO-3306, 2 µM C-604) (Fig. 3e). B55α depletion did not mitigate the mitotic defects induced by partial CDK1 inhibition (siB55α; 2 µM RO-3306, 0 µM C-604), indicating that these defects arose directly from compromised CDK1 function rather than PP2A-B55α derepression (Fig. 3e). In HeLa cells, B55α depletion under conditions of strong CDK1 inhibition (siB55α; 4 µM RO-3306, 0 µM C-604) increased the incidence of mitotic slippage and skipping while decreasing the frequency of mitotic catastrophe (Fig. 3e). This shift was less evident in RPE-1 cells, where treatment with 4 µM RO-3306 predominantly induced mitotic catastrophe, irrespective of B55α expression (siB55α; 4 µM RO-3306, 0 µM C-604) (Fig. 3e). Together, these data demonstrated how PP2A-B55α activity shapes mitotic pathologies caused by CDK1 and GWL inhibition and highlighted cell-type-specific differences in reliance on PP2A-B55α activity during mitotic exit.

### Multivariate analysis identifies PP2A-B55α-dependent G2 regulation as the primary point of cell-type-specific divergence

While combined inhibition of CDK1 and GWL produced phenotypes broadly consistent across the two cell lines, reduced PP2A-B55α activity caused markedly different, cell-type-specific effects on both interphase control and mitotic fidelity. These differences suggested marked variation in how distinct cell types integrate PP2A-B55α into cell-cycle control and motivated further characterisation of the most pronounced divergences. To rank divergent responses in an unbiased manner, we partitioned the dataset into four groups defined by cell line and transfected siRNA: (1) HeLa siCTR, (2) HeLa siB55α, (3) RPE-1 siCTR and (4) RPE-1 siB55α. We then used principal component analysis (PCA) to project group-specific interphase and mitotic responses across six combinations of RO-3306 (0, 2, 4 µM) and C-604 (0, 2 µM) treatments into a two-dimensional space (Fig. 3f, Materials and Methods). Within this space, we calculated group-wise Euclidean distances between the projected phenotypic outputs and converted them to z-scores (z) by standardising across all computed values (Fig. 3g). Among the resulting 36 distances, G2-specific responses of B55α-depleted HeLa cells showed the greatest divergence, highlighting the role of PP2A-B55α in G2/M control as the key discrepancy between tumour-derived HeLa and non-transformed RPE-1 cells (Fig. 3g).

### Modelling the cancer-associated signature characterised by elevated GWL and reduced B55α expression

The cell-type-specific role of PP2A-B55α in G2/M regulation was of particular interest, given that a subset of tumours (13.2%) showed relatively low B55α and high GWL expression, a pattern absent from tumour-adjacent normal tissues (Fig. 1c, e and Supplementary Fig. S1). Since dysregulation of PP2A-B55α and GWL has been implicated in cancer ^49–52^, we sought to model this putative cancer signature across different cellular systems and assess its impact on fundamental cellular functions and vulnerabilities. To overexpress GWL, we established stable HeLa (WT and PIP-FUCCI), RPE-1, and HCC1395 (triple-negative breast cancer, providing a second cancer-derived model) cell lines harbouring an integrated, doxycycline-inducible, Flag-tagged GWL construct (GWL-OE), with peak expression ∼24 h after doxycycline administration (Supplementary Fig. S6a, b). Overexpression of the GWL-Flag protein rescued the effects of C-604-mediated GWL inhibition, confirming that the construct remained enzymatically active and that the GWL-OE system could be used to assess the consequences of varying GWL activity levels (Supplementary Fig. S6c-g). To complete the expression signature of interest, we combined the GWL-OE system with siRNA-mediated depletion of B55α. Although B55α depletion was equally efficient across cell lines and growth conditions, GWL overexpression was more pronounced in B55α-depleted cells, indicating that PP2A-B55α may limit GWL levels via an unknown mechanism (Fig. 4a, b and Supplementary Fig. S7a-c). Nevertheless, the established system qualitatively modelled the GWL-dominant imbalance between B55α and GWL expression and enabled characterisation of its functional consequences in three distinct cellular contexts.

**Fig. 4.**
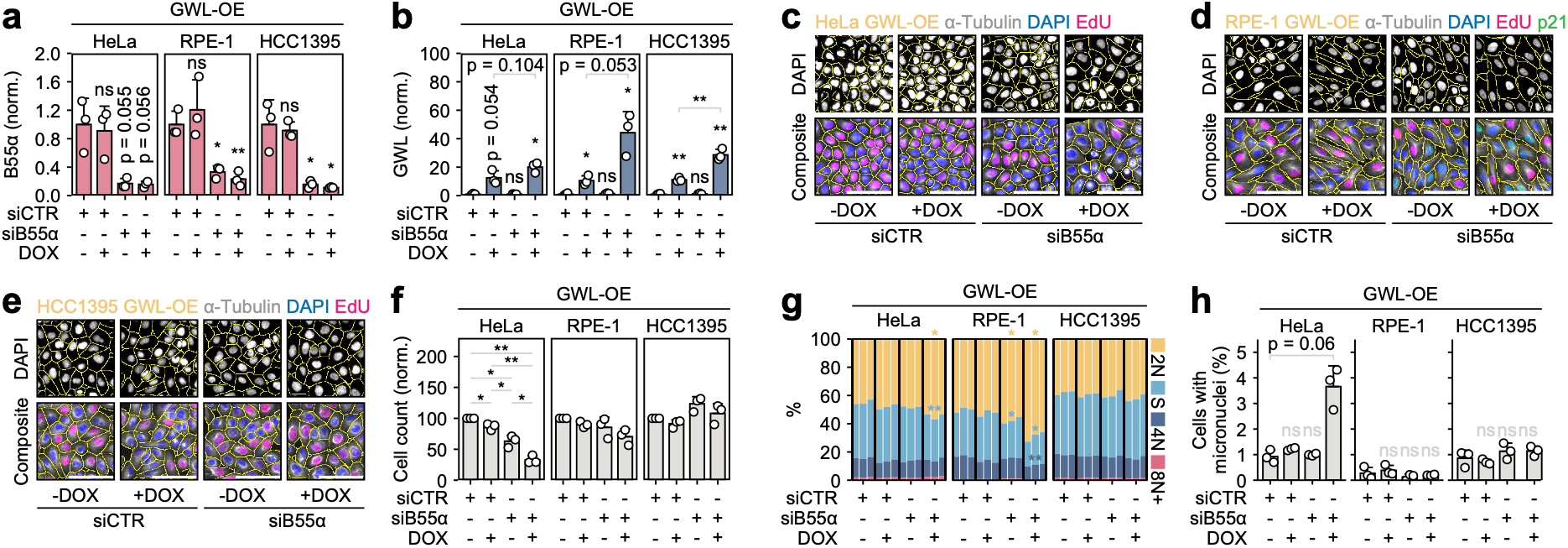
Modelling of the recurrent cancer-associated signature characterised by the relative dominance of GWL over B55α. **(a-b)** Quantification of B55α (a) and GWL (b) protein levels based on western blots from (Supplementary Fig. S7a-c) in HeLa, RPE-1 and HCC1395 GWL-OE cells. Densitometric values were normalised to the loading control and to the mean of the (siCTR, −DOX) control condition. Bars and error bars represent means and standard deviations across n = 3 biological replicates, with individual values shown as points. Statistical significance of relevant comparisons was assessed using Welch’s two-tailed t-tests, without p-value correction. ns – not significant, * p < 0.05, ** p < 0.01, *** p < 0.001. For panels (C-H), HeLa, RPE-1, and HCC1395 GWL-OE cells were analysed across four perturbation conditions: control siRNA (siCTR) or B55α-targeting siRNA (siB55α), each grown without (−DOX) or with (+DOX) doxycycline. Analyses presented in (C-H) are derived from the same imaging dataset. **(c-e)** Representative images of HeLa (c), RPE-1 (d) and HCC1395 (e) GWL-OE cells under the indicated perturbation conditions. Composite (α-tubulin, DAPI, EdU, and p21 in RPE-1 GWL-OE cells only) and single-channel (DAPI) images are shown. Cellular segmentation is indicated by yellow outlines. Scale bars are 100 µm. **(f)** Proliferation represented by total cell count (based on cell segmentation) from immunofluorescence imaging. Bars and error bars represent means and standard deviations across n = 3 biological replicates, with individual values shown as points. Statistical significance was assessed using Welch’s two-tailed t-tests for all pairwise comparisons between conditions within each cell line, with Benjamini-Hochberg correction across the six pairwise tests per cell line. ns – not significant, * p < 0.05, ** p < 0.01. **(g)** Proportions of cell-cycle groups, classified as 2N (G1), S (EdU-positive), 4N (G2/M), and polyploid (8N+), derived from imaging-based DAPI integrated intensity and EdU incorporation. For each condition, the proportions of cell-cycle phases are shown as three stacked bars, representing results from n = 3 biological replicates. Statistical significance of phase-specific proportion differences between each treatment condition and the (siCTR, −DOX) control was assessed using Welch’s two-tailed t-tests, with p-values corrected using the Benjamini-Hochberg procedure. Only significant comparisons are indicated. * p < 0.05, ** p < 0.01. **(h)** Manually determined frequencies of cells with micronuclei. Bars and error bars represent means and standard deviations, respectively. Points represent results from n = 3 biological replicates. In each experiment, between 671 and 1925 cells were scored per condition. Statistical significance was assessed using Welch’s two-tailed t-tests comparing each condition with the (siCTR, −DOX) control, with p-values corrected using the Benjamini-Hochberg procedure. ns – not significant.

The long-term effects of combined B55α depletion and GWL overexpression (120 h post-siRNA transfection, 96 h post-doxycycline) varied across three cell lines. The most severe chronic phenotypes were observed in HeLa cells (siB55α, +DOX), which exhibited a pronounced proliferation defect, an accumulation of 2N cells, increased cell size, and a higher frequency of micronuclei, consistent with increased chromosomal instability (Fig. 4c, f-h and Supplementary Fig. S7d-f). The response in RPE-1 cells (siB55α, +DOX) was more moderate, characterised by a subtle proliferation defect, a higher proportion of 2N cells, an insignificant increase in cell size, and no detectable increase in micronuclei formation (Fig. 4d, f-h and Supplementary Fig. S7d-f). Following B55α depletion, p53-proficient RPE-1 cells also displayed a moderate upregulation of p21 (Supplementary Fig. S7g-i). Although this p21 accumulation was not further potentiated by GWL overexpression, it likely contributed to the observed accumulation of 2N cells and mild proliferation defect (Supplementary Fig. S7g-i). In contrast to HeLa and RPE-1 cells, HCC1395 cells (siB55α, +DOX) remained largely unaffected, maintaining normal growth and cell-cycle distribution without morphological abnormalities or a higher incidence of micronuclei (Fig. 4e-h and Supplementary Fig. S7d-f). Notably, even in the most affected HeLa cells, a combination of low B55α and high GWL expression levels did not abolish proliferation, as HeLa GWL-OE cells (siB55α, +DOX) formed macroscopic colonies, albeit of reduced size (Supplementary Fig. S7j, k). These findings further indicated that the cellular consequences of PP2A-B55α perturbations depend on cellular context and raised questions about the mechanisms that shape them.

### B55α depletion and GWL overexpression increase sensitivity to inhibition of WEE1 and PKMYT1 in tumour-derived cells

Given the role of the GWL/PP2A-B55α module in G2/M control, we hypothesised that the cellular consequences of its perturbation are, at least in part, shaped by the activity of the CDK1-inhibitory kinases WEE1 and PKMYT1. To test this hypothesis, we assessed how HeLa, RPE-1, and HCC1395 GWL-OE cells responded to increasing doses of the WEE1 inhibitor adavosertib ^65^ and the PKMYT1 inhibitor RP-6306 ^66^. In non-transformed RPE-1 GWL-OE cells, B55α depletion and GWL overexpression did not increase sensitivity to either inhibitor, indicating either minimal functional overlap between PP2A-B55α and WEE1/PKMYT1 or the involvement of other G2/M regulators that compensate for reduced PP2A-B55α activity (Fig. 5a-c and Supplementary Fig. S8, a-c). In tumour-derived HeLa and HCC1395 GWL-OE cells, B55α depletion and GWL overexpression synergistically increased the cytostatic potential of both adavosertib and RP-6306, suggesting that, in these cells, PP2A-B55α cooperated with both WEE1 and PKMYT1 (Fig. 5a-c and Supplementary Fig. S8a-c). Notably, the functional overlap between PP2A-B55α and PKMYT1 appeared stronger than that between PP2A-B55α and WEE1, as sensitisation to RP-6306 (siB55α +DOX ED_50_ fold-change ± SD; HeLa: 18.6 ± 7.4, HCC1395: 5.7 ± 1.6) exceeded that to adavosertib (HeLa: 6.3 ± 1.4, HCC1395: 1.8 ± 0.1) (Fig. 5a-c and Supplementary Fig. S8a-c).

**Fig. 5.**
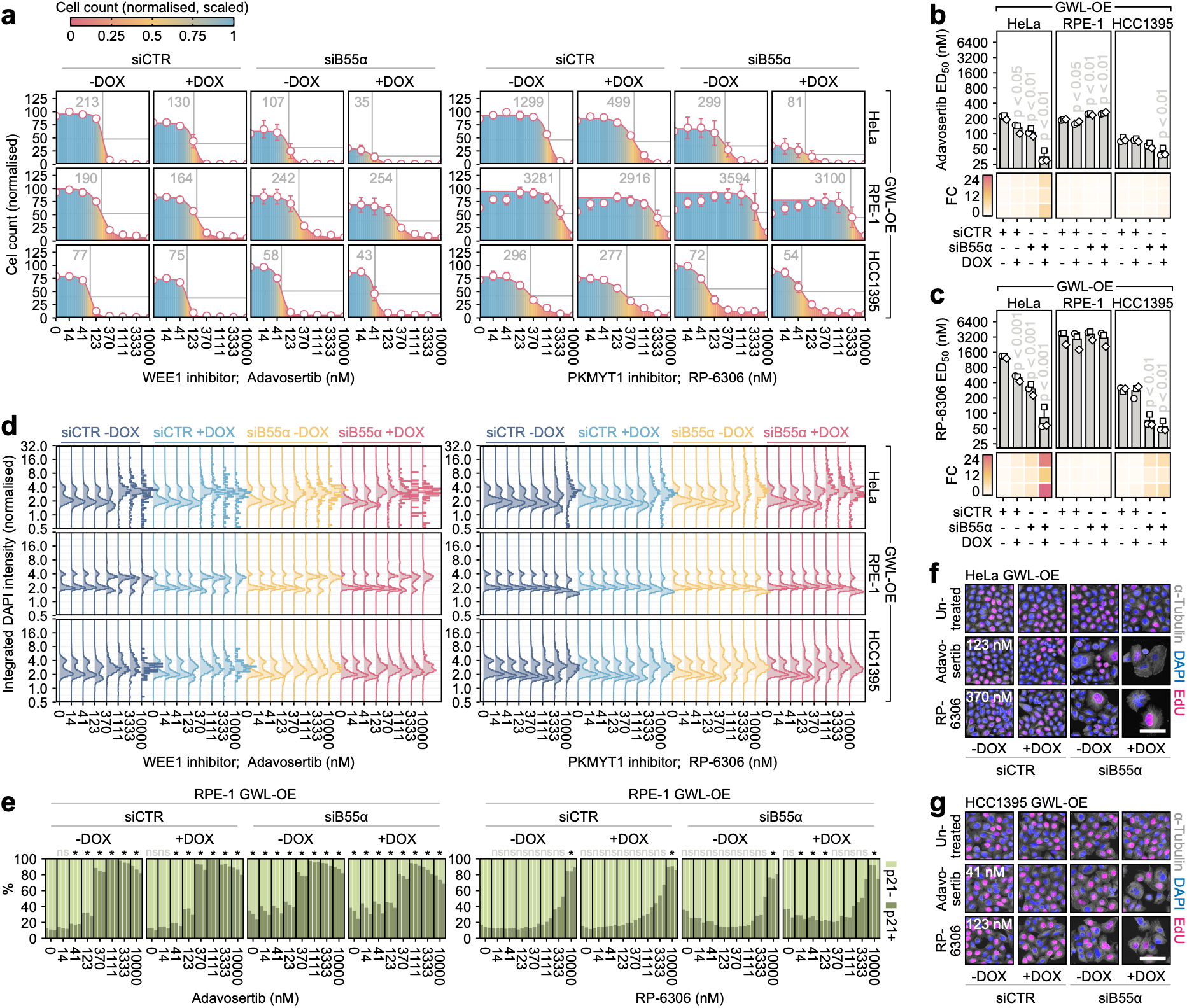
Cell-type-specific effects of low B55α and high GWL expression on cellular responses to inhibition of WEE1 and PKMYT1. **(a)** Dose-response curves showing the effects of the WEE1 inhibitor adavosertib and the PKMYT1 inhibitor RP-6306 on HeLa, RPE-1, and HCC1395 GWL-OE cells across the four perturbation conditions. Cytotoxicity was assessed as total cell count (cell segmentation) from immunofluorescence imaging. Cell counts are normalised to the maximum value within each cell line, preserving baseline differences between conditions arising from the cytotoxic effects of B55α depletion and GWL overexpression. Points and error bars represent means and standard deviations across n = 3 biological replicates. Lines represent the mean of three independently fitted four-parameter sigmoidal models. The overlay colour gradients show the same data renormalised within each condition (each curve scaled from 0 to 1), allowing comparison of drug-response shapes independent of baseline differences. Mean ED_50_ values are indicated. **(b, c)** Cellular sensitivities to Adavosertib (b) and RP-6306 (c). Top panels: ED_50_ estimates from each of n = 3 biological replicates (points), with bars and error bars showing means and standard deviations, respectively. Statistical significance of comparisons between each condition and the (siCTR, −DOX) control was assessed using Welch’s two-tailed t-tests, with p-values corrected using the Benjamini-Hochberg procedure. Bottom panels: heatmaps showing the fold change in ED_50_ relative to the (siCTR, −DOX) control; each row represents one biological replicate. **(d)** Ridgeline plots showing dose-dependent effects of adavosertib and RP-6306 on DNA content distributions in HeLa, RPE-1, and HCC1395 GWL-OE cells under the four perturbation conditions, measured as integrated DAPI intensity per cell from immunofluorescence imaging data. Each ridge represents pooled single-cell measurements across n = 3 biological replicates. Counts varied with treatment: HeLa, 29-94,521 cells per condition (median 50,338); RPE-1, 2,542-59,505 (median 37,084); HCC1395, 84-48,495 (median 33,706). **(e)** Proportions of p21-positive (p21+) and p21-negative (p21-) RPE-1 GWL-OE cells across the four perturbation conditions, treated with adavosertib (0-10 µM) or RP-6306 (0-10 µM). For each condition, the proportions of p21+/p21-cells are shown as three stacked bars, representing results from n = 3 biological replicates. Statistical significance of comparisons between each condition and the untreated (siCTR, −DOX) control was assessed using Welch’s two-tailed t-tests, with Benjamini-Hochberg correction across the comparisons. Significance is shown qualitatively. ns – not significant, * p < 0.05. **(f, g)** Representative immunofluorescence images (composite; α-tubulin, DAPI, EdU) of HeLa (f) and HCC1395 (g) GWL-OE cells under the four perturbation conditions, treated with adavosertib or RP-6306 at doses that showed the strongest differential effects: 123 nM adavosertib and 370 nM RP-6306 for HeLa; 41 nM adavosertib and 123 nM RP-6306 for HCC1395. The presented panels represent subsets of the larger panels shown in (Supplementary Fig. S8a-c). Scale bars, 100 µm.

In RPE-1 GWL-OE cells, adavosertib (≥370 nM) induced the accumulation of enlarged 4N cells and upregulated p21 expression, consistent with cell-cycle arrest. In contrast, RP-6306 (10,000 nM) caused minor shifts in DNA content, accompanied by high p21 expression, but only a limited increase in cell size (Fig. 5d, e and Supplementary Fig. S8b). In HeLa and HCC1395 GWL-OE cells, adavosertib and RP-6306 induced marked, dose-dependent shifts in DNA content distributions and significant cellular and nuclear pathologies, likely reflecting defective mitotic progression (Fig. 5d, f, g and Supplementary Fig. S8, c). The dose-dependent occurrence of these defects was exacerbated by B55α depletion and GWL overexpression, suggesting that, in HeLa and HCC1395 GWL-OE cells, impaired PP2A-B55α activity synergised with WEE1 or PKMYT1 inhibition by compromising mitotic control (Fig. 5d, f, g and Supplementary Fig. S8a, c). Despite this synergy, expression levels of B55α and GWL, alone or in combination, did not predict cellular sensitivity to either adavosertib or RP-6306 across a minimal panel of six cell lines, all of which lacked established markers of WEE1 or PKMYT1 dependence (Supplementary Figs. S9 and S10). Together, these findings indicated that dependence on WEE1 and PKMYT1 is shaped by multiple factors, including additional, as-yet uncharacterised determinants.

### A combination of defective p53 signalling and broader PP2A-B55 perturbation creates PKMYT1 dependency in non-transformed RPE-1 cells

The lack of synergy between reduced PP2A-B55α activity and inhibition of WEE1 or PKMYT1 in RPE-1 cells suggested that this cell line employed compensatory mechanisms to offset compromised PP2A-B55α function. Two plausible candidates were intact p53 signalling, which is defective in both HeLa and HCC1395 cells, and the deployment of PP2A-B55δ, an alternative PP2A-B55 holoenzyme incorporating the B55δ regulatory subunit. Individually, p53 knockout or siRNA-mediated depletion of B55α and B55δ had only limited effects on the dose-dependent cytotoxicity of either adavosertib or RP-6306 (Fig. 6a-e). However, when combined, these three perturbations produced marginal sensitisation to adavosertib (AUC FC ± SD = 0.79 ± 0.12) and moderate sensitisation to RP-6306 (AUC FC ± SD = 0.58 ± 0.03) (Fig. 6a-f). These results indicated that cellular dependence on PKMYT1 can emerge through the cumulative disruption of multiple cellular functions that redundantly reinforce the G2/M boundary. Additionally, the differential sensitisation to adavosertib and RP-6306 further suggested that PKMYT1 and WEE1 occupy distinct regulatory niches, with PKMYT1 aligning more closely with PP2A-B55 complexes.

**Fig. 6.**
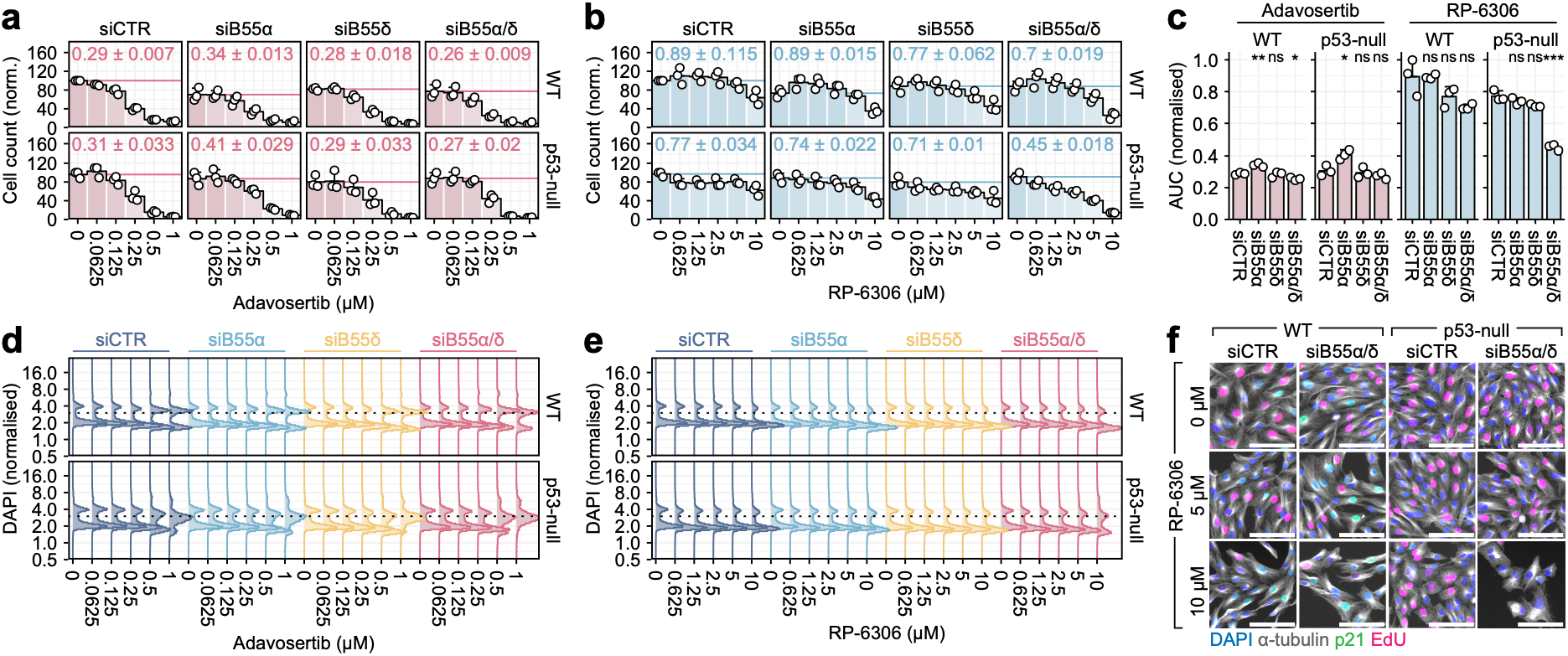
The effects of p53 deficiency and broader PP2A-B55 perturbation on PKMYT1 and WEE1 dependency in non-transformed RPE-1 cells. **(a-b)** The effect of increasing doses of adavosertib (a) and RP-6306 (b) on the proliferation of RPE-1 WT and RPE-1 p53-null cells transfected with siRNAs targeting B55α or B55δ. Proliferation is represented by total cell counts inferred from immunofluorescence imaging. Cell counts are normalised to the maximum value within each cell line. Bars and error bars represent means and standard deviations across n = 3 biological replicates. Points represent the results of individual experiments. Mean AUC values, normalised to untreated control, and their standard deviations are indicated. (c) Cellular sensitivities to adavosertib and RP-6306, expressed as normalised AUC values. Bars and error bars are means and standard deviations across n = 3 biological replicates. Points represent the results of individual experiments. Statistical significance of differences between each condition and the control (siCTR) was assessed by an unpaired two-tailed t-test with Benjamini-Hochberg correction. ns – not significant, * p < 0.05, ** p < 0.01, *** p < 0.001. (d-e) Ridgeline plots showing dose-dependent effects of adavosertib (d) and RP-6306 (e) on DNA content distributions in RPE-1 WT and RPE-1 p53-null cells under the indicated conditions, measured as integrated DAPI intensity per cell from immunofluorescence imaging data. Each ridge represents pooled single-cell measurements across n = 3 biological replicates. (f) Immunofluorescence images (composite; α-tubulin, DAPI, p21, EdU) of RPE-1 WT and p53-null cells transfected with both siB55α and siB55δ and treated with 5 or 10 µM RP-6306.

### Sensitisation to WEE1 and PKMYT1 inhibition operates via distinct mechanisms

Although WEE1 and PKMYT1 kinases are generally considered functionally redundant in restraining mitotic entry through inhibitory phosphorylation of CDK1, perturbation of PP2A-B55α in HeLa and HCC1395 cells produced markedly stronger sensitisation to the PKMYT1-selective RP-6306 than to the WEE1-selective adavosertib. This discrepancy suggested that WEE1 and PKMYT1 are not fully redundant and that they form distinct functional relationships with PP2A-B55α. To explore the basis of these relationships, we examined the effects of B55α depletion and GWL overexpression on S- and G2-phase progression and mitotic fidelity in HeLa PIP-FUCCI GWL-OE cells treated with increasing doses of adavosertib or RP-6306 (Supplementary movies M4 and M5). Consistent with the roles of WEE1 and PKMYT1 in preventing premature mitotic commitment, cytostatic doses of both inhibitors induced analogous mitotic failures, which occurred during the first or second cell division after drug administration (siCTR −DOX; ED_50_ ± SD: adavosertib 0.25 ± 0.02 µM, RP-6306 2.55 ± 0.46 µM) (Fig. 7a-d and Supplementary Fig. S11a, b). Consistent with its uneven impact on the cytostatic effects of both inhibitors, depletion of B55α increased cellular susceptibility to mitotic defects induced by adavosertib and RP-6306 by 1.9- and 3.7-fold, respectively (siB55α −DOX; ED_50_ ± SD: adavosertib, 0.13 ± 0.03 µM; RP-6306, 0.69 ± 0.12 µM) (Fig. 7b-d and Supplementary Fig. S11a, b). These uneven effects were further exacerbated by GWL overexpression, which, in combination with B55α depletion, amplified sensitivity to adavosertib by 2.5-fold and to RP-6306 by 8.2-fold (siB55α +DOX; ED_50_ ± SD: adavosertib 0.10 ± 0.02 µM, RP-6306 0.31 ± 0.13 µM) (Fig. 7b-d and Supplementary Fig. S11a-b). Thus, although adavosertib and RP-6306 induced apparently analogous mitotic defects, reduced PP2A-B55α activity sensitised cells to the two inhibitors to markedly different extents.

**Fig. 7.**
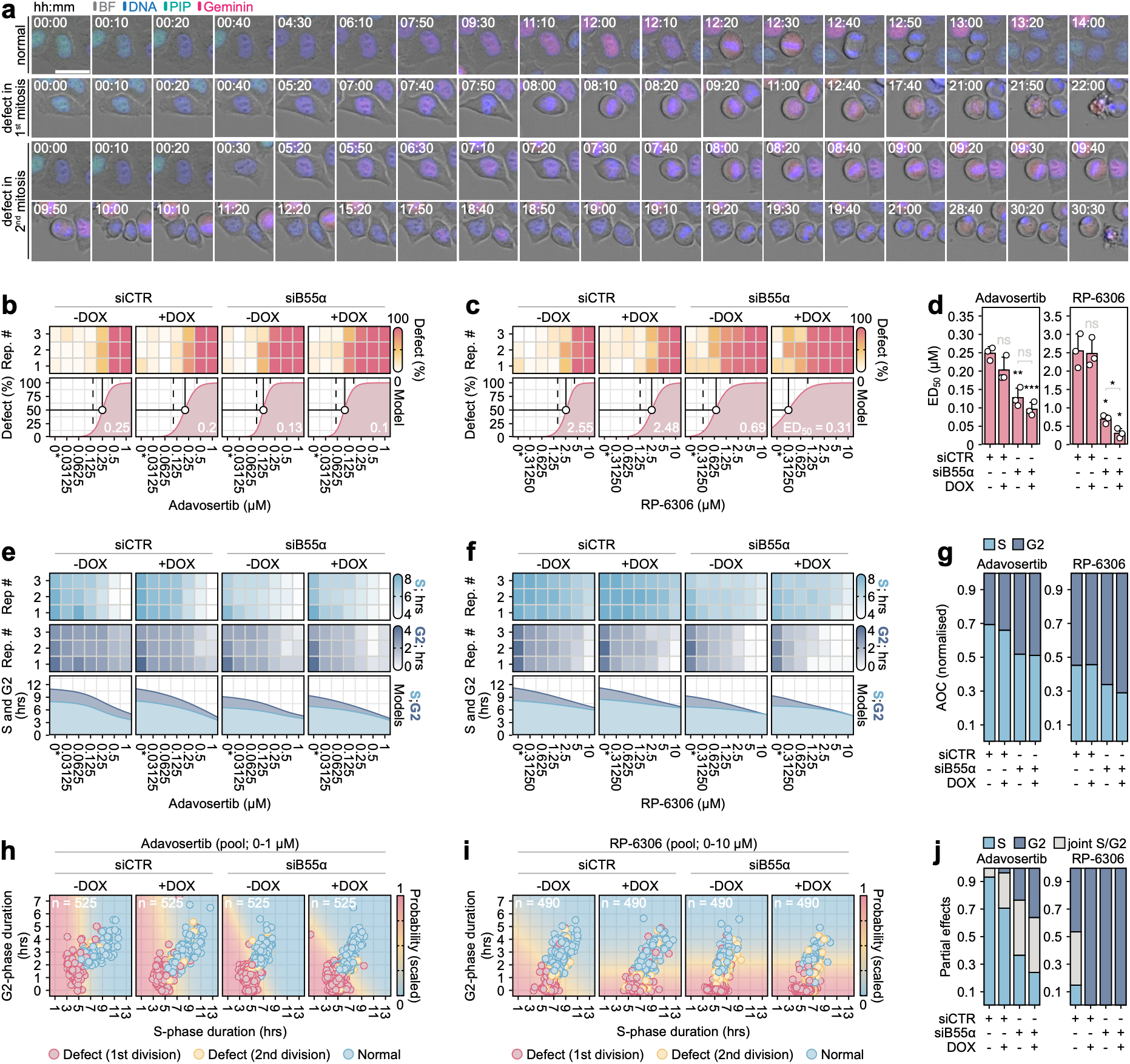
Synergy between compromised PP2A-B55α activity and inhibition of WEE1 or PKMYT1. **(a)** Representative time-lapse images (composite: bright field, DNA, PIP-eGFP, Geminin-mCherry) of live HeLa PIP-FUCCI GWL-OE cells treated with cytotoxic doses of adavosertib or RP-6306, illustrating normal and defective mitotic outcomes in the first or second division after drug administration. Time stamps are shown. Scale bar, 30 µm. **(b-c)** Dose-dependent frequencies of mitotic defects induced by adavosertib (b; 0-1 µM) and RP-6306 (c; 0-10 µM) in HeLa PIP-FUCCI GWL-OE cells across the four perturbation conditions (siCTR or siB55α, each with or without DOX). Top panels: heatmaps of mitotic defect frequencies (0-100%), with each row representing one of n = 3 biological replicates (20-25 mitotic events were scored per condition per replicate). Bottom panels: means of three fitted four-parameter sigmoidal models. Mean ED_50_ doses are indicated numerically and by a solid vertical line; ED_10_ and ED_90_ doses are marked by vertical dashed lines. Zero-dose (0*) is plotted at 0 on the heatmap and at half the lowest dose on the model panel for a log_2_-axis display. **(d)** Adavosertib and RP-6306 ED_50_ estimates from each of n = 3 biological replicates (points), with bars and error bars showing means and standard deviations, respectively. Statistical significance of comparisons between each condition and the (siCTR, −DOX) control, and between (siB55α −DOX) and (siB55α +DOX), was assessed using Welch’s two-tailed t-tests, with p-values corrected using the Benjamini-Hochberg procedure across the four comparisons per drug. ns – not significant, * p < 0.05, ** p < 0.01, *** p < 0.001. **(e-f)** Dose-dependent changes in S- and G2-phase durations induced by adavosertib (e; 0-1 µM) and RP-6306 (f; 0-10 µM) in HeLa PIP-FUCCI GWL-OE cells across the four perturbation conditions. Top panels: heatmaps showing median S- and G2-phase durations, with each row representing one of n = 3 biological replicates (20-25 single cells were analysed per condition per replicate). Bottom panels: means of three fitted four-parameter sigmoidal models for S-and G2-phase durations, displayed as stacked area plots. Zero-dose (0*) is plotted at 0 on the heatmaps and at half the lowest dose on the model panels for log-axis display. **(g)** Normalised Area Over Curve (AOC) values quantifying the dose-dependent shortening of S- and G2-phase durations, derived from (e-f). For each phase, AOC was computed as the trapezoidal integral of (maximum observed duration − dose-specific duration) across log_2_-transformed dose, capturing the cumulative shortening relative to the maximum observed value. AOC values were normalised within each condition so that the sum of the S- and G2-phase integrals equals 1, representing the proportional contribution of each phase to the total response. **(h-i)** Logistic regression models predicting mitotic defect outcome from S- and G2-phase durations in HeLa PIP-FUCCI GWL-OE cells across the four perturbation conditions, treated with adavosertib (h) or RP-6306 (i). Cells from all doses (adavosertib: 0-1 µM; RP-6306: 0-10 µM) and n = 3 biological replicates were pooled per condition (adavosertib: 525 cells, RP-6306: 490 cells), capturing the full range of S/G2 durations induced across the increasing doses. Each point represents a single cell, coloured by observed outcome (normal, defect at first division, or defect at second division). Background colour shows the probability of mitotic defect predicted by elastic net logistic regression (binomial family, α = 0.75; λ selected via cross-validation at the λ.1SE value). Predictions were evaluated on a uniform grid across the displayed S and G2 ranges, with observed cells (points) indicating the data range used for model fitting. **(j)** Variance decomposition of the logistic regression linear predictors from (h, i), showing the proportional contributions of S-phase duration, G2-phase duration, and their joint covariance (S/G2) to the variance of model predictions across the four perturbation conditions. Contributions were computed from the regression coefficients and predictor (co)variances (see Methods) and sum to 1.

To assess the attributes of interphase events preceding normal and defective mitoses, we quantified the durations of the preceding S and G2 phases, capturing both single-dose effects and responses integrated across the dose ranges (quantified as phase-specific areas above the curves; AOCs) (Fig. 7e-g and Supplementary Fig. S11c-h). Both adavosertib and RP-6306 shortened S- and G2-phase durations in a dose-dependent manner, but with distinct phase biases. Adavosertib predominantly reduced S-phase duration, whereas RP-6306 primarily affected the G2 phase (Fig. 7e-g and Supplementary Fig. S11c-h). Under both treatment regimes, B55α depletion and GWL overexpression had limited impact on dose-dependent S-phase effects but exacerbated G2-phase shortening (Fig. 7e-g and Supplementary Fig. S11c-h). Notably, this G2-specific synergy was modest in adavosertib-treated cells but pronounced following RP-6306 treatment, indicating that PP2A-B55α cooperates more closely with PKMYT1 than with WEE1 in regulating the G2/M transition. This suggested that uneven sensitisation to adavosertib and RP-6306 in cells with compromised PP2A-B55α function may reflect the phase-separated functions of WEE1 and PKMYT1 in S- and G2-phase control, respectively.

To test this hypothesis at single-cell resolution, we used elastic-net logistic regression models to predict the probability of mitotic failure from S- and G2-phase durations. We fitted eight models, one for each combination of siRNA (siCTR or siB55α), doxycycline (−DOX or +DOX), and treatment (0-1 µM adavosertib or 0-10 µM RP-6306), enabling us to compare the predictive contributions of S- and G2-phase durations across distinct perturbations (Fig. 7h, i). By decomposing the variance of the linear predictor, we partitioned predictions of mitotic failure into S- and G2-specific coefficients and a joint S/G2 coefficient reflecting the covariance between S- and G2-phase durations (coordinated shortening of S and G2) (Fig. 7j). In cells with unperturbed B55α and GWL levels, prediction of adavosertib-induced mitotic defects was driven mainly by S-phase length, with minimal G2-specific and joint contributions (Fig. 7j). This prediction shifted progressively towards G2 and joint S/G2 coefficients following B55α depletion or GWL overexpression, with the most evident redistribution observed when both perturbations were combined (Fig. 7j). In contrast, the prediction of RP-6306-induced mitotic failure in otherwise unperturbed cells was primarily driven by joint S/G2 and G2 coefficients, with minimal contribution from S-phase durations (Fig. 7j). Notably, after B55α depletion or GWL overexpression, S and joint S/G2 coefficients were reduced to zero, leaving G2-phase durations as the sole predictors (Fig. 7j). Together, these results supported a model in which compromised PP2A-B55α function destabilises the G2/M boundary and creates a G2-specific reliance on PKMYT1 to prevent premature G2/M transition and consequent mitotic failure.

### Tumours preserve relatively normal WEE1 expression but upregulate PKMYT1

The functional divergence between WEE1 and PKMYT1 was further supported by their distinct cancer-associated expression patterns. According to our TCGA analysis, tumours rarely dysregulated WEE1 expression but frequently upregulated PKMYT1, indicating that these two CDK1-inhibitory kinases are subject to distinct selective pressures during tumour evolution (Fig. 8a, b). Notably, normal tissues generally exhibited higher WEE1 than PKMYT1 expression, whereas tumours often displayed a reciprocal pattern, with PKMYT1 levels exceeding those of WEE1 (Fig. 8b). Although the physiological consequences of this expression pattern remain to be established, PKMYT1 overexpression was already evident in Stage I tumours across a broad range of cancer types, suggesting that increased PKMYT1 activity may promote early tumour development (Fig. 8c). Given the frequent perturbations of the CDK1/GWL/PP2A-B55α axis and other mitotic entry regulators, PKMYT1 overexpression may represent an adaptive mechanism that stabilises the G2/M boundary and supports the proliferation of cellular states with compromised control of mitotic commitment. Defining the molecular foundations of these states will be essential for understanding how tumour cells remodel G2/M control networks and for developing robust predictive procedures to identify patients most likely to benefit from PKMYT1-targeted therapy.

**Fig. 8.**
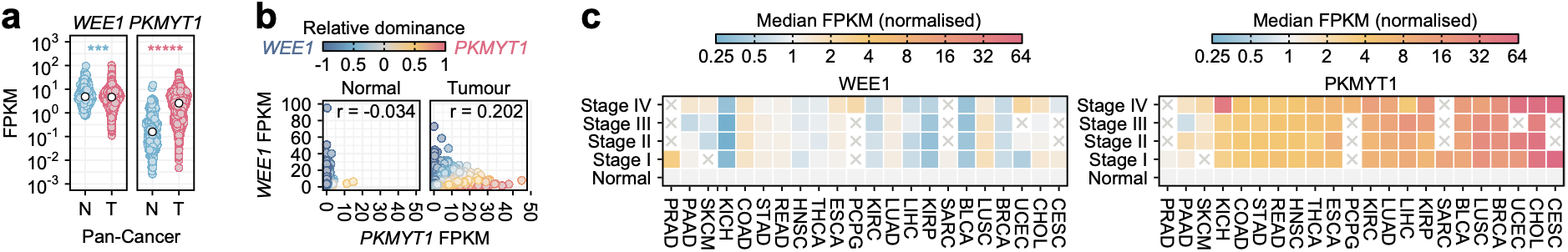
Cancer-associated expression profiles of WEE1 and PKMYT1. **(a)** WEE1 and PKMYT1 expression from TCGA data in tumour-adjacent normal (N; 23 tissue types, n = 723) and tumour (T; 33 cancer types; n = 10,296) tissues, shown as violin point plots. Coloured points represent individual samples, and white points represent the medians. Statistical significance was assessed using the Wilcoxon rank-sum test with Benjamini-Hochberg correction. *** p < 0.001, ***** p < 0.00001. Asterisks are colour-coded to indicate the direction of the difference. **(b)** Scatterplot showing RNA expression levels (FPKM) of *WEE1* and *PKMYT1* across TCGA pan-cancer samples shown in (A). Colour gradients represent relative dominance (RD) scores, calculated as RD = (x - y) / (x + y), where x and y are the gene expression values on the x- and y-axes, respectively. r – Pearson correlation coefficient. **(c)** Heatmaps showing median WEE1 and PKMYT1 FPKM values across stage-stratified cancers with at least n = 3 matching normal tissues (n = 22). In each cancer group, FPKM medians are normalised to the respective normal tissue values. BLCA – bladder urothelial carcinoma, BRCA – breast invasive carcinoma, CESC – cervical squamous cell carcinoma and endocervical adenocarcinoma, CHOL – cholangiocarcinoma, COAD – colon adenocarcinoma, ESCA – oesophageal carcinoma, HNSC – head and neck squamous cell carcinoma, KICH – kidney chromophobe, KIRC – kidney renal clear cell carcinoma, KIRP – kidney renal papillary cell carcinoma, LIHC – liver hepatocellular carcinoma, LUAD – lung adenocarcinoma, LUSC – Lung squamous cell carcinoma, PAAD – pancreatic adenocarcinoma, PCPG – pheochromocytoma and paraganglioma, PRAD – prostate adenocarcinoma, READ – rectum adenocarcinoma, SARC – sarcoma, SKCM – skin cutaneous melanoma, STAD – stomach adenocarcinoma, THCA – thyroid carcinoma, UCEC – uterine corpus endometrial carcinoma

## Discussion

The CDK1/GWL/PP2A-B55α signalling axis is frequently dysregulated in cancer, and perturbations of its components are implicated in genomic instability, defective checkpoint control, uncontrolled proliferation, and metastasis. Tumour tissues frequently exhibit multiple simultaneous perturbations of this axis, yet the interactions among these alterations and their collective impact on cell-cycle regulation remain poorly understood. Here, we systematically characterise the effects of combined perturbations of CDK1, GWL, and PP2A-B55α on interphase control, mitotic progression, and cellular dependence on the CDK1-inhibitory kinases WEE1 and PKMYT1.

Consistent with the mutual reinforcement between CDK1 and GWL activation during the G2/M transition, reduced CDK1 activity and near-complete GWL inhibition synergistically increased G2 duration in both HeLa and RPE-1 cells. Combined and single-agent effects of CDK1 and GWL inhibition were attenuated by B55α depletion, confirming a conserved role of PP2A-B55α in reinforcing the G2/M boundary under conditions of reduced CDK1 and/or GWL activity. GWL inhibition alone, however, caused only marginal G2 prolongation, in contrast to observations in *Xenopus* egg extracts, where GWL-mediated inhibition of PP2A-B55δ is essential for mitotic entry ^19,20^. This discrepancy suggests that, in human cells, GWL primarily stabilises phosphorylation of mitotic substrates once the point of mitotic commitment is reached and, consistent with its proposed role in checkpoint recovery ^67^, contributes to G2/M timing predominantly when CDK1 activity is limiting.

Although the CDK1 activation threshold is widely regarded as a barrier that ensures timely initiation and successful completion of mitosis ^1,2^, our findings suggest that cells can enter mitosis even when CDK1 activity is insufficient to complete it. We show that reducing effective CDK1 activity leads to stage-specific mitotic failures, ranging from late mitotic exit defects, such as micronuclei and incomplete cytokinesis, to pre-anaphase mitotic catastrophe, early mitotic slippage, and complete mitotic skipping. Consistent with the proposed model that pre-anaphase mitotic events are driven by progressive CDK1 activation ^4,63^, these results suggest that transitions between mitotic stages occur at discrete thresholds of CDK1 activity. Indeed, graded CDK1 perturbations produce the same spectrum of mitotic outcomes in both tumour-derived HeLa and non-transformed RPE-1 cells, indicating that distinct thresholds of CDK1-dependent substrate phosphorylation are a general mitotic control mechanism rather than a phenomenon specific to cancer cells. These findings may point towards a refinement of the mitotic bistable switch hypothesis, in which CDK1 activity could be more precisely tuned at different stages of mitosis than previously appreciated. Multiple research groups have resolved phosphoproteomes of distinct mitotic stages, including prophase, prometaphase, metaphase, and mitotic exit; however, causal links between specific phosphorylation events and phase transitions remain elusive ^24,26,31,68,69^. Ultimately, why cells enter mitosis with suboptimal CDK1 activity remains a critical question, particularly given that cellular stress responses typically converge on CDK1 inhibition to prevent premature mitotic entry ^70,71^. Characterising the molecular determinants of mitotic CDK1 activity thresholds and how they integrate into the broader G2/M control network will therefore be essential to understanding their contributions to normal and pathological cellular functions.

Our findings provide new insights into the architecture of G2/M control and reveal that the functional organisation of its components varies across cell types. In tumour-derived HeLa cells, PP2A-B55α plays a broader role in reinforcing the G2/M boundary, and reducing its activity accelerates mitotic entry even under otherwise unperturbed conditions. This function appears to be absent in non-transformed RPE-1 cells, which maintain normal G2/M timing independently of PP2A-B55α. While it remains unclear why different cell types adopt distinct configurations of G2/M control, our data suggest that reorganising the mitotic entry network can create exploitable vulnerabilities. The cell-type-specific role of PP2A-B55α in G2/M control overlaps with that of PKMYT1, and compromised PP2A-B55α activity, achieved through B55α depletion or GWL overexpression, sensitises HeLa and breast cancer-derived HCC1395 cells to the PKMYT1 inhibitor RP-6306. By contrast, RPE-1 cells, which do not rely on PP2A-B55α to reinforce the G2/M boundary, remain tolerant to PKMYT1 inhibition regardless of B55α or GWL status. As 13.2% of TCGA tumour cases exhibit an expression pattern characterised by relatively low B55α levels and concurrent GWL overexpression, this finding may be clinically relevant. However, our analysis also indicates that B55α and GWL expression levels alone do not reliably predict RP-6306 toxicity, suggesting the involvement of other contributing factors. These may include other G2/M regulators or endogenous sources of replication stress, which have emerged as key determinants of PKMYT1 dependency (CITATION). Consistent with the more complex view of PKMYT1 dependence, co-depletion of B55α and its paralogue B55δ in p53-deficient RPE-1 cells produced moderate sensitisation to PKMYT1 inhibition, indicating that reliance on PKMYT1 can be exacerbated by a broader disruption of PP2A-B55 function, combined with defective p53-dependent checkpoint signalling.

Interestingly, reduced PP2A-B55α activity had only limited effects on cellular sensitivity to the WEE1 inhibitor adavosertib. Thus, despite the shared ability of PKMYT1 and WEE1 to restrain CDK1 activity, PP2A-B55α dysfunction selectively increases cellular dependence on PKMYT1. While this selective relationship could, at least in part, reflect additional PKMYT1 functions in chromosome segregation ^72^ or non-canonical substrates beyond CDK1(T14) ^73^, our analysis indicates that it primarily arises from the distinct contributions of PKMYT1 and WEE1 to interphase control. Logistic modelling of S- and G2-phase lengths and their impact on mitotic progression revealed that mitotic failures induced by PKMYT1 inhibition are predominantly associated with shortened G2, whereas those induced by WEE1 inhibition are more closely linked to S-phase shortening and show only a weak association with G2 duration. These differences are consistent with the broader substrate specificity of nuclear WEE1, which regulates both CDK2 and CDK1 during the S and G2 phases ^5,6,74–76^, and with the selective function of PKMYT1, which inhibits cytoplasmic CDK1 and primarily controls G2 progression ^7,8^. Together, these findings indicate that, in certain contexts, PP2A-B55α and PKMYT1 converge to restrain CDK1 activity during G2 via complementary mechanisms, with PP2A-B55α reversing CDK1-dependent phosphorylation and PKMYT1 directly inhibiting CDK1. These complementary functions likely ensure sufficient G2 duration to complete synthesis of late-replicating loci and prevent premature mitotic entry with unresolved replication intermediates. Failure of this safeguard may generate under-replicated DNA that, as described following WEE1 inhibition, becomes susceptible to MUS81-SLX4-mediated processing, resulting in DNA breaks, chromosome pulverisation and mitotic cell death ^75–77^. However, whether mitotic death induced by PKMYT1 inhibition involves replication-associated lesions, particularly when G2/M control is disrupted, remains to be determined.

Further supporting functional divergence between the two CDK1-inhibitory kinases, our analysis revealed that WEE1 expression in tumour tissues remains largely unchanged, whereas PKMYT1 is frequently upregulated. We speculate that this discrepancy reflects their distinct positions within cell-cycle control. While WEE1 occupies a broader niche by controlling both CDK2 and CDK1, potentially imposing stronger constraints on its dysregulation, PKMYT1 provides a more selective mechanism to reinforce G2/M control without directly affecting S-phase progression. In theory, such selective reinforcement may be particularly important in cancer cells with compromised thresholds for mitotic entry and a higher propensity to initiate the cell division programme in the presence of under-replicated DNA.

So far, PKMYT1 dependence has been linked to CCNE1 amplification, loss-of-function mutations in FBXW7 (which encodes an SCF E3 ubiquitin ligase component that targets cyclin E1 for degradation), and deleterious mutations in PPP2R1A, the Aα scaffolding subunit of PP2A complexes ^78–80^. While CCNE1 amplification and FBXW7 loss promote PKMYT1 dependence through cyclin E1 accumulation and replication stress, the consequences of PPP2R1A alterations remain less well understood. Despite these mechanistic associations, overexpression of CCNE1 and deleterious FBXW7 or PPP2R1A mutations did not reliably predict responses to single-agent PKMYT1 inhibition in a biomarker-selected oncology cohort, suggesting that these features alone are insufficient to define PKMYT1 dependency ^78,80^. While we propose B55α depletion and GWL overexpression as additional context-dependent determinants of cellular reliance on PKMYT1, we also note that they likely do not satisfy the criteria for reliable predictive biomarkers. Broadly, our work proposes high PKMYT1 dependence as a joint consequence of diverse perturbations that compromise G2/M control and increase the probability of mitotic entry with under-replicated DNA, a vulnerability that is likely to be particularly pronounced in cells experiencing replication stress. Ultimately, defining the combinations of alterations that generate this vulnerable state, together with the compensatory mechanisms that stabilise it, will be critical for predicting which tumours are most likely to benefit from PKMYT1-targeted therapy.

## Methods

### Cell culture

All cell lines were cultured at 37 °C in a humidified atmosphere with 5% CO2. HeLa and U2OS cells were maintained in Dulbecco’s modified Eagle medium (DMEM; Gibco, 21969035); hTERT RPE-1 cells in DMEM/F-12 (Sigma-Aldrich, D8437); and HCC1395, HCC1143 and BT-549 cells in RPMI-1640 (Gibco, 31870025). All media were supplemented with penicillin-streptomycin (Sigma-Aldrich, P4333) and either 10% fetal bovine serum (Gibco, A5670701) or 10% tetracycline-free fetal bovine serum (Pan Biosciences, P30-3602) for doxycycline-inducible lines. DMEM and RPMI-1640 were also supplemented with 2 mM L-glutamine (Sigma-Aldrich, G7513). For live-cell imaging, HeLa and hTERT RPE-1 cells were cultured in FluoroBrite DMEM (Gibco, A1896701) and DMEM/F-12 without phenol red (Gibco, 11039021), respectively. All cell lines were routinely tested for mycoplasma contamination.

### Generation of PIP-FUCCI cell lines

HeLa and RPE-1 PIP-FUCCI cell lines were generated by lentiviral transduction. Lentiviral particles were produced in HEK293T cells by co-transfection with the viral envelope protein vector pCMV-VSV-G (Addgene, 8454), the 2^nd^-generation lentiviral packaging vector psPAX2 (Addgene, 12260), and the PIP-FUCCI expression vector pLenti-PGK-Neo-PIP-FUCCI (Addgene, 118616). Transfection was performed using the calcium/phosphate mammalian transfection kit (Clontech Laboratories, 631312). The transfection mixture (2.5 µg of total plasmid DNA and 124 mM calcium phosphate in 100 µL of HEPES-buffered saline) was incubated for 10 min at 25 °C and then added dropwise to 50-80% confluent HEK293T cells. Following transfection, cells were incubated overnight at 37 °C (5% CO_2_). The original medium was replaced with fresh medium, and the cells were incubated for an additional 24 h. To harvest the viral particles, the medium was collected, centrifuged (1,200 × g, 3 min, 25 °C), and filtered through a 0.45 µm filter. Recipient HeLa and RPE-1 cells were seeded at 30-40% confluence 24 h before infection. The filtered viral supernatant was supplemented with 8 µg/mL polybrene and added to the recipient cells. 24 hours after transduction, the original medium was replaced with fresh medium containing G418 (500 µg/mL). PIP-FUCCI-positive cells were single-cell sorted using the BD FACSMelody Cell Sorter system and cultured for 3-4 weeks in the presence of G418 (500 µg/mL). Selected clones were expanded and validated by immunofluorescence microscopy, assessing DNA content (integrated DAPI intensity) and cellular morphology (α-tubulin staining), and by live-cell imaging to validate expression of PIP-FUCCI probes.

### Generation of GWL overexpression cell lines

HeLa, HeLa PIP-FUCCI, RPE-1 and HCC1395 GWL-OE cell lines were generated using the Sleeping Beauty (SB) transposase system ^81^ with two plasmids: an SB transposase plasmid (Addgene, 34879) and a derivative of an SB transposon plasmid carrying a blue fluorescent protein (BFP) marker (Addgene, 60521) with the FLAG-tagged GWL construct under a doxycycline-inducible promoter. Plasmids were transfected using Lipofectamine LTX with PLUS reagent (Thermo Fisher Scientific, A12621). To generate HeLa, HeLa PIP-FUCCI and HCC1395 GWL-OE cell lines, 70,000 HeLa and HeLa PIP-FUCCI cells and 120,000 HCC1395 cells were seeded in a 12-well plate. The following day, the original medium was removed and replaced with 0.9 mL of fresh medium. 100 µL of the transfection mixture (component A: 125 ng of SB transposase, 250 ng of SB transposon, 1.2 µL of PLUS reagent, up to 50 µL of MEM; component B: 47 µL of MEM, 3 µL of Lipofectamine LTX) was then added dropwise. To generate the RPE-1 GWL-OE cell line, 140,000 RPE-1 cells were seeded into a 6-well plate. The following day, the original medium was removed and replaced with 1.9 mL of fresh medium. 100 µL of the transfection mixture (component A: 26 ng of SB transposase, 474 ng of SB transposon, 1.2 µL of PLUS reagent, up to 50 µL of MEM; component B: 47 µL of MEM, 3 µL of Lipofectamine LTX) was then added dropwise. Transfection mixture components were prepared separately, incubated for 5 min at 25 °C, then combined and incubated for an additional 25 min before addition. Seven days after transfection, BFP-positive cells were single-cell sorted using the BD FACSMelody Cell Sorter system and cultured for 3-4 weeks. Single clones were validated by immunofluorescence microscopy for DNA content (integrated DAPI intensity), cellular morphology (α-tubulin staining), and GWL-Flag inducibility (anti-FLAG or anti-GWL staining after doxycycline treatment).

### siRNA transfection

Cells at 70-90% confluence were harvested by trypsinisation, counted with a haemocytometer, and resuspended at 1.07 × 10^5^ cells/mL. A total of 1.8 mL of cell suspension (1.93 × 10^5^ cells) was mixed with siRNA transfection mix containing 600 µL of MEM medium (Gibco, 21090022), 6 µL of Lipofectamine RNAiMAX (Thermo Fisher Scientific, 13778100), and 2.4 µL of 20 µM siRNA SMARTpool (Dharmacon), yielding a final siRNA concentration of 20 nM. Transfection mixes were pre-incubated at 25 °C for 30 min before addition. The following SMARTpools were used: non-targeting control (siCTR; D-001810-10-20), PPP2R2A/B55α (siB55α; L-004824-00-0005), and PPP2R2D/B55δ (siB55δ; L-032298-00-0005). Transfected cells were incubated for 24 h, then trypsinised and reseeded into the appropriate experimental format (60-mm dishes or 96-well plates).

### Colony formation

Cells were harvested by trypsinisation, counted with a haemocytometer, and diluted to 80 cells/mL. Three mL (240 cells) were seeded into each 60-mm dish (Corning, 430166). Dishes were incubated at 25 °C for 20-30 min, then transferred to 37 °C (5% CO_2_) for 5-6 h to allow cell attachment. Subsequently, 1 mL of fresh medium containing either DMSO or the drug of interest (at 4× the final concentration) was added, yielding a final volume of 4 mL. After 72 h, the medium was replaced with 4 mL of drug-free medium, and dishes were incubated for 9-18 days, depending on the cell line. Colonies were fixed with 3.7% formaldehyde containing 1% methanol (Fisher Scientific, 10160052) in PBS at 25 °C for 15 min and stained with 0.01% crystal violet (Sigma-Aldrich, V5265) in PBS for 30 min. Excess crystal violet was washed off with distilled H_2_O. Stained dishes were scanned using an office scanner (Epson), and colonies were quantified using Cellpose (version 2.0.5) with custom-trained segmentation models specific to each cell line.

### Protein extraction and Western blotting

Cells were harvested by trypsinisation or mitotic shake-off and, depending on the size of the cell pellet, resuspended in 20-100 µL of 9 M urea supplemented with 5% 2-mercaptoethanol. Cell lysates were sonicated and incubated at 95 °C for 5 min. Protein concentrations were determined using the Qubit 4 system (Invitrogen) with the Qubit protein assay kit (Invitrogen, 33211). Protein samples were diluted and mixed with 4× Laemmli sample buffer (Bio-Rad, 1610747) to achieve a final concentration of 1 µg/µL. 10-20 µg of protein were resolved on a 10% or 12% SDS-polyacrylamide gel and transferred onto a PVDF membrane (Amersham, 10600023) using the Trans-Blot Turbo Transfer System (Bio-Rad) at a constant 25 V for 30 min. The transfer used a three-buffer system with the following layers: three sheets of filter paper soaked in anode 1 buffer (300 mM Tris, 20% methanol, pH 10.4), two sheets of filter paper soaked in anode 2 buffer (25 mM Tris, 20% methanol, pH 10.4), a membrane soaked in methanol, the gel containing size-resolved proteins, and five sheets of filter paper soaked in cathode buffer (25 mM Tris, 40 mM 6-aminohexanoic acid, 20% methanol, pH 9.4). The membrane with transferred proteins was washed with methanol for 30 sec and then submerged in distilled H_2_O. The membrane was stained with 0.1% (w/v) Ponceau S (Sigma-Aldrich, P3504) in 5% acetic acid and imaged to visualise the transfer. The membrane was de-stained by several washes in distilled H_2_O and blocked with 3% BSA in TBST (20 mM Tris, 150 mM NaCl, 0.1% Tween 20, pH 7.6) for 60 min at 25 °C. The membrane was probed with primary antibodies diluted in 3% BSA in TBST at 4 °C overnight. The membrane was washed three times with TBST (5, 5, 15 min) and then probed with HRP-conjugated secondary antibodies for 60 min at 25 °C. The membrane was washed three times with TBST (5, 5, 15 min) and covered with the Clarity Western ECL substrate (Bio-Rad, 1705061). For targets with low signal, this was supplemented with 10% (v/v) SuperSignal West Femto Maximum Sensitivity Substrate (Thermo Scientific, 34095). The membrane was imaged using the ImageQuant LAS 4000 (GE Healthcare) system, and protein band intensities were quantified by densitometry in ImageJ (2.16.0) ^82^. The following primary antibodies were used: anti-GWL raised in rabbit (1:1,000; Sigma-Aldrich, HPA027175), anti-PRC1(T481)-P raised in rabbit (1:1,000; Abcam, ab62366), anit-B55α raised in rabbit (1:1,000; CST, 2290S), anti-GAPDH raised in mouse (1:20,000; Abcam, ab8245), anti-β-Actin raised in mouse (1:5,000, Abcam, ab3280), anti-histone H3 raised in rabbit (1:5,000, CST, 4499S). The following secondary antibodies were used: anti-mouse HRP-conjugated, raised in goat (1:5,000; Dako, P0447), and anti-rabbit HRP-conjugated, raised in goat (1:2,000; Dako, P0448).

### Immunofluorescence imaging

Immunofluorescence experiments were performed in black 96-well plates with optically clear, flat bottoms (Revvity, 6055302). For a 96-hour incubation, the following seeding densities were used: 750 cells/well (HeLa), 1,000 cells/well (RPE-1) and 1,000 cells/well (HCC1395). For experiments involving S-phase labelling, cells were treated with 5-ethynyl-2’-deoxyuridine (EdU; 10 µM) for 20 min before fixation. Cells were fixed with methanol-free 4% formaldehyde (CST, 47746) for 10 min at 25 °C, then washed with 100 µL of PBS. Fixed cells were permeabilised and blocked with blocking solution (CST, 12411) for 60 min at 25 °C, then washed with 100 µL of PBS. If the experiment involved EdU labelling, cells were incubated with 50 µL of click-iT reaction mixture containing 43 µL of TBS (50 mM Tris-HCl, 150 mM NaCl, pH 7.4), 2 µL of 100 mM CuSO_4_, 0.12 µL of 0.1 mg/mL Sulfo-Cy5-Azide (Jena Bioscience, CLK-AZ118-5) and 5 µL of 1.12 M (+)-Sodium L-ascorbate (Sigma-Aldrich, 11140) for 30 min at 25 °C, then washed three times with 100 µL of PBS. Cells were covered with 50 µL of antibody dilution buffer (CST, 12378) containing primary antibodies and incubated at 4 °C overnight. Cells were washed three times with 100 µL of PBS, covered with 50 µL of antibody dilution buffer containing fluorescently labelled secondary antibodies and incubated for 30 min at 25 °C. Cells were washed three times with 100 µL of PBS, covered with 50 µL of 1 µg/mL 4′,6-diamidino-2-phenylindole (DAPI) and incubated for 30 min at 25 °C. Cells were washed once with 100 µL of PBS and covered with 200 µL of PBS. Plates were stored at 4 °C and imaged within 14 days. The following primary antibodies were used: anti-α-tubulin raised in mouse (1:1,000; CST, 3873); anti-α-tubulin raised in rat (1:1000; Novus Biologicals, NB600-506); anti-p21 raised in rabbit (1:800; CST, 2947); anti-GWL raised in rabbit (1:1,000; Sigma-Aldrich, HPA027175); and anti-FLAG raised in mouse (1:1,000; Sigma-Aldrich, F1804). The following secondary antibodies were used: anti-mouse Alexa Fluor 488 conjugate raised in goat (1:2,000; CST, 4408S); anti-rabbit Alexa Fluor 555 conjugate raised in goat (1:2,000; CST, 4413S); anti-mouse Alexa Fluor 546 conjugate raised in goat (1:2,000; Thermo Fisher Scientific, A11003); anti-rabbit Alexa Fluor 647 conjugate raised in donkey (1:2,000; Thermo Fisher Scientific, A-31573); and anti-rat Alexa Fluor 488 conjugate raised in donkey (1:2,000; Thermo Fisher Scientific, A21208). Plates were imaged using the Operetta CLS high-content system (Revvity).

### Live-cell imaging

Live cells were imaged in black 96-well plates with optically clear, flat bottoms (Revvity, 6055302). 24 hours before imaging, cells were seeded in 300 µL of phenol-red-free medium supplemented with 1:5,000 SPY650-DNA (Spirochrome, SC501) at seeding densities of 2,000 cells/well (HeLa PIP-FUCCI), 2,000 cells/well (HeLa PIP-FUCCI GWL-OE), and 3,000 cells/well (RPE-1 PIP-FUCCI). Before imaging, the medium was replaced with fresh, pre-tempered phenol-red-free medium containing SPY650-DNA. To minimise potential stress, cells were allowed to recover for 4-8 hours at 37 °C (5% CO_2_). Cells were imaged using the pre-equilibrated (37 °C, 5% CO_2_) Operetta CLS high-content analysis system (Revvity) with a 20× air objective (NA = 0.4, WD = 8.28). Four channels were imaged at each time point: brightfield (BF), green (EGFP, PIP-eGFP), red (mCherry; Geminin-mCherry), and far-red (Alexa 647; SPY650). HeLa and RPE-1 PIP-FUCCI cells were imaged at 10- and 15-min intervals, respectively.

### Processing and analysis of immunofluorescence imaging data

Raw immunofluorescence imaging data were stored in the OMERO cloud repository ^83^. Cellular and nuclear masks were generated with Cellpose ^84^ (version 3.1.1.1) using custom, cell-type-specific segmentation models trained on α-tubulin and DAPI images, respectively. Mean and integrated fluorescence intensities were quantified using a custom analysis pipeline (https://github.com/HocheggerLab/omero-screen). Quantified intensities were normalised to the values under the first peak in the respective intensity distributions under control conditions. This normalisation was also applied to integrated DAPI intensities, which indicated relative DNA content (2N, 4N, 8N+). Downstream single-cell analysis was performed in R (version 4.2.3).

### Processing and analysis of live-cell imaging data

Raw live-cell movies were stored in the OMERO cloud repository ^83^ and processed using custom-built computational pipelines (https://github.com/HocheggerLab/gttrack.git). Individual images were flatfield-corrected, stitched, and segmented with Cellpose (version 3.1.1.1) ^84^ together with custom segmentation models to generate nuclear masks. Time-resolved stitched nuclear masks were tracked using Trackastra (version 0.2.3) ^85^, and resulting lineages were stored in Cell Tracking Challenge (CTC) format. A subset of tracked lineages was manually curated using the Mastodon ImageJ plugin (version 1.0.0) to ensure accuracy. For interphase analysis, each curated track was initiated at cell birth and terminated at the first frame showing mitotic entry (Geminin-mCherry signal migrating from the nucleus to the cytoplasm) or mitotic skipping (loss of the Geminin-mCherry signal). Curated nuclear masks were used to quantify time-resolved mean nuclear PIP-eGFP and Geminin-mCherry fluorescence intensities, which were further analysed in R (version 4.2.3). Briefly, PIP-eGFP and Geminin-mCherry intensity tracks were smoothed with a rolling mean (9-time-point window) and normalised using min-max scaling. Normalised tracks were combined into a difference index: (Geminin − PIP) / (Geminin + PIP), scaled between -1 and 1, where Geminin and PIP represent the smoothed normalised intensities of PIP-eGFP and Geminin-mCherry, respectively. The difference index was smoothed (9-time-point rolling mean) and converted to moving differences calculated as x(i+1) - x(i). Moving differences were smoothed (9-time-point rolling mean) and used to segment each track into G1, S, and G2 phases. The G1/S transition was defined as the latest time point preceding the peak of the moving difference that has a moving difference score < 25% of the peak value. The S/G2 transition was defined as the earliest time point after the peak of the moving difference with a moving difference score > 125% of the minimum value recorded between the peak and the end of the track. In cases of mitotic skipping, where manual track termination was inaccurate, tails showing declining Geminin-mCherry signal (< 90% of the maximum value) were trimmed.

### Statistical analysis

All analyses were conducted in R (4.2.3) using the rstatix (0.7.2), FSA (0.10.0), emmeans (1.11.1), lme4 (2.0.1), drc (3.0.1), pracma (2.4.4) and glmnet (4.1.8) packages. Per-replicate analyses (n = 3 biological replicates) used Welch’s two-tailed t-tests by default, with Benjamini-Hochberg correction applied across pre-specified comparison families. For comparisons of multiple treatment conditions against a single reference control (used in panels assessing perturbation effects on mitotic-outcome proportions), per-replicate values were analysed using a linear model with treatment group as a fixed effect, followed by Dunnett’s post hoc test. Validation experiments addressing biologically distinct hypotheses were tested without multiple-comparison correction, as noted in the corresponding legends. The standard significance threshold scheme was applied: * p < 0.05, ** p < 0.01, *** p < 0.001, **** p < 0.0001. Pooled single-cell analyses used the Kruskal-Wallis test followed by Dunn’s post hoc test with Bonferroni correction. Given the large effective sample sizes, a stricter threshold scheme was applied: * p < 0.01, ** p < 0.001, *** p < 0.0001, **** p < 0.00001. TCGA pan-cancer analyses (tumour-adjacent normal, n = 723; tumour, n = 10,296) used two-sided Wilcoxon rank-sum tests, with the stricter threshold scheme applied throughout. Dose-response curves were fitted with four-parameter sigmoidal models. For RO-3306 and C-604 inhibitor characterisation, a single model was fitted to all replicates within each condition. For adavosertib and RP-6306 dose-response analyses, models were fitted to each biological replicate independently. Area Under the Curve (AUC) and Area Over the Curve (AOC) values were computed by trapezoidal integration over the log-transformed dose axis. For multivariate analyses, principal component analysis (PCA) was performed on centred and scaled response vectors. Pairwise Euclidean distances in PCA space were z-scored using the combined distribution. Logistic regression modelling of mitotic outcomes used elastic net regularisation (binomial family, α = 0.75), with the regularisation parameter λ selected via cross-validation using the λ.1SE criterion.

### AI usage declaration

Artificial intelligence tools, including Grammarly (1.175.0.0), Claude (Opus 4.7), and ChatGPT (5.5), were used to refine, correct, edit, and format the text to improve clarity and accuracy of language. Claude (Opus 4.7) was used to aid the code development for trapezoidal integration analyses and logistic regression analysis. Claude (Opus 4.7) was used to scrutinise statistical analysis, including the use of appropriate statistical tests and their reporting.

## Data availability

Raw data can be obtained from the following repositories:

https://doi.org/10.6084/m9.figshare.33046862 (Colony formation analysis; CF),

https://doi.org/10.6084/m9.figshare.33047003 (Immunofluorescence analysis; IF),

https://doi.org/10.6084/m9.figshare.33047030 (Live-cell analysis; LC)

https://doi.org/10.6084/m9.figshare.33047048 (Western blot analysis; WB)

https://doi.org/10.6084/m9.figshare.33047075 (TCGA analysis)

https://doi.org/10.6084/m9.figshare.33047117 (DepMap analysis)

## Code availability

https://github.com/HocheggerLab/omero-screen (Image processing and quantification)

https://github.com/HocheggerLab/gttrack.git (Live-cell imaging tracking

https://github.com/R-Zach/Perturbation-of-the-CDK1-GWL-PP2A-B55-axis-reveals-PP2A-B55-as-a-cell-type-specific-G2-M-regulator.git (Other analyses)

## Acknowledgments

This work was supported by funding from Cancer Research UK – C28206/A14499 (R.Z., A.D.H., M.M., L.G., W.R.F., H.H.) and Wellcome Trust – 110578/Z/15/Z (H.H.).

## Author contributions

Conceptualisation: R.Z., H.H.

Experimental design: R.Z., H.H.

Experiments: R.Z., L.G.

Dasta analysis: R.Z., H.H.

Methods development: R.Z., A.D.H., M.M., L.G., W.R.F., H.H.

Supervision: R.Z., H.H.

Writing original draft: R.Z., H.H.

Consultation and review: C.M.D.

R.Z. conceptualised the research project, designed and conducted experiments, analysed and visualised the data, and wrote the manuscript; A.D.H. developed code to process IF and live-cell imaging data; C.M.D. provided consultation and reviewed the manuscript; M.M. generated stable PIP-FUCCI cell lines; L.G. contributed to optimising live-cell experiments and to establishing the tracking-correction procedures; W.R.F. established the GWL overexpression system; H.H. conceptualised the project, designed experiments, developed code to process IF imaging data, and wrote the manuscript.

## Supplementary Information

## Supplementary Figures

**Supplementary Fig. S1.**
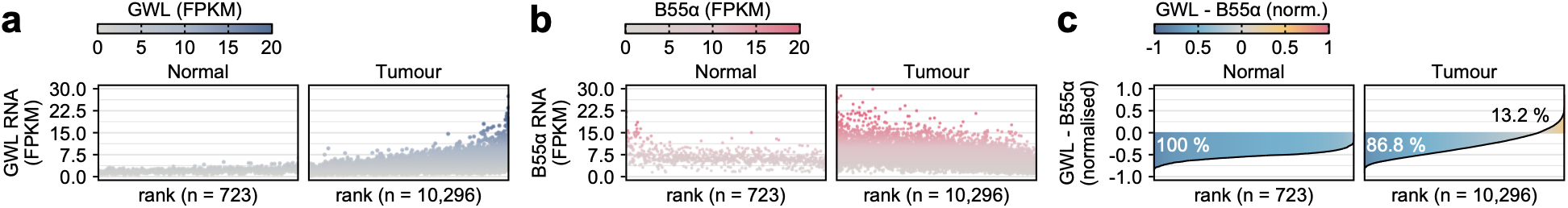
Recurrent cancer signature defined by relatively low B55α and high GWL expression. **(a, b)** Ranked RNA expression levels (FPKM) of GWL (a) and B55α (b) in TCGA pan-cancer tumour-adjacent normal (n = 723) and tumour (n = 10,296) samples, ordered by the relative dominance (RD) scores shown in (C). **(c)** Ranked RD scores, calculated as (GWL − B55α) / (GWL + B55α), as defined in Figure 1. Positive values indicate GWL-dominant samples; negative values indicate B55α-dominant samples. Proportions of samples in each category are shown.

**Supplementary Fig. S2.**
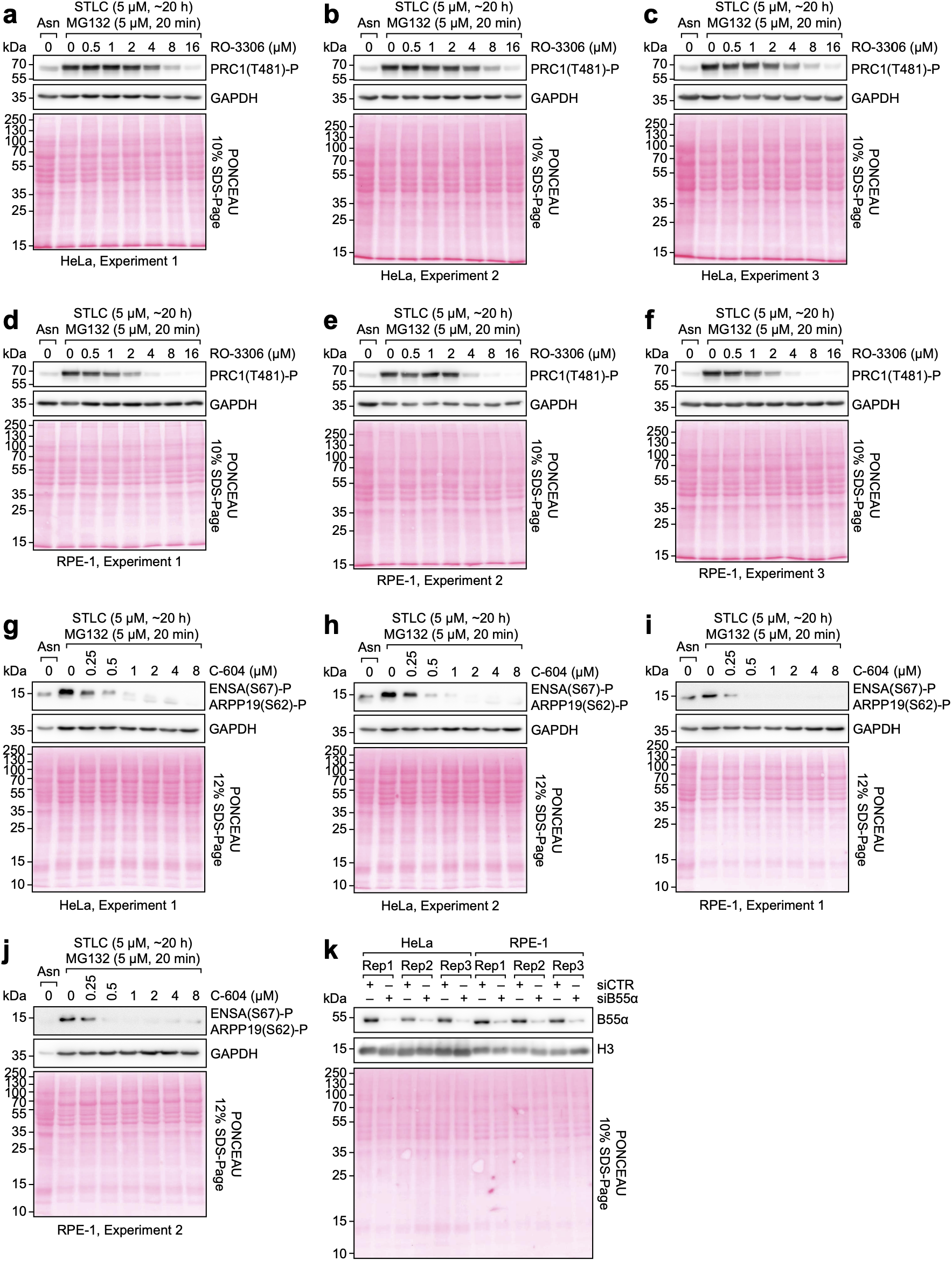
Perturbations of CDK1, GWL and PP2A-B55α. **(a-f)** Western blots showing dose-dependent effects of RO-3306 on PRC1 phosphorylation at threonine 481 (T481) in mitotic HeLa (a-c; n = 3 biological replicates) and RPE-1 (d-f; n = 3 biological replicates) cells. **(g-j)** Western blots showing dose-dependent effects of C-604 on phosphorylation of ENSA at serine 67 (S67) and ARPP19 at serine 62 (S62) in mitotic HeLa (g, h; n = 2 biological replicates) and RPE-1 (i, j; n = 2 biological replicates) cells. (a-j) Cells were arrested in prometaphase with S-trityl-L-cysteine (STLC; 5 µM) for 20 hours, then treated with the indicated concentrations of RO-3306 or C-604 in the presence of STLC and the proteasome inhibitor MG132 (5 µM) for 20 minutes. GAPDH served as a loading control. **(k)** Representative western blot showing the effect of B55α-targeting siRNA (siB55α) compared to control (siCTR) on B55α protein levels in asynchronous HeLa and RPE-1 cells, 48 hours post-transfection. Histone H3 served as a loading control.

**Supplementary Fig. S3.**
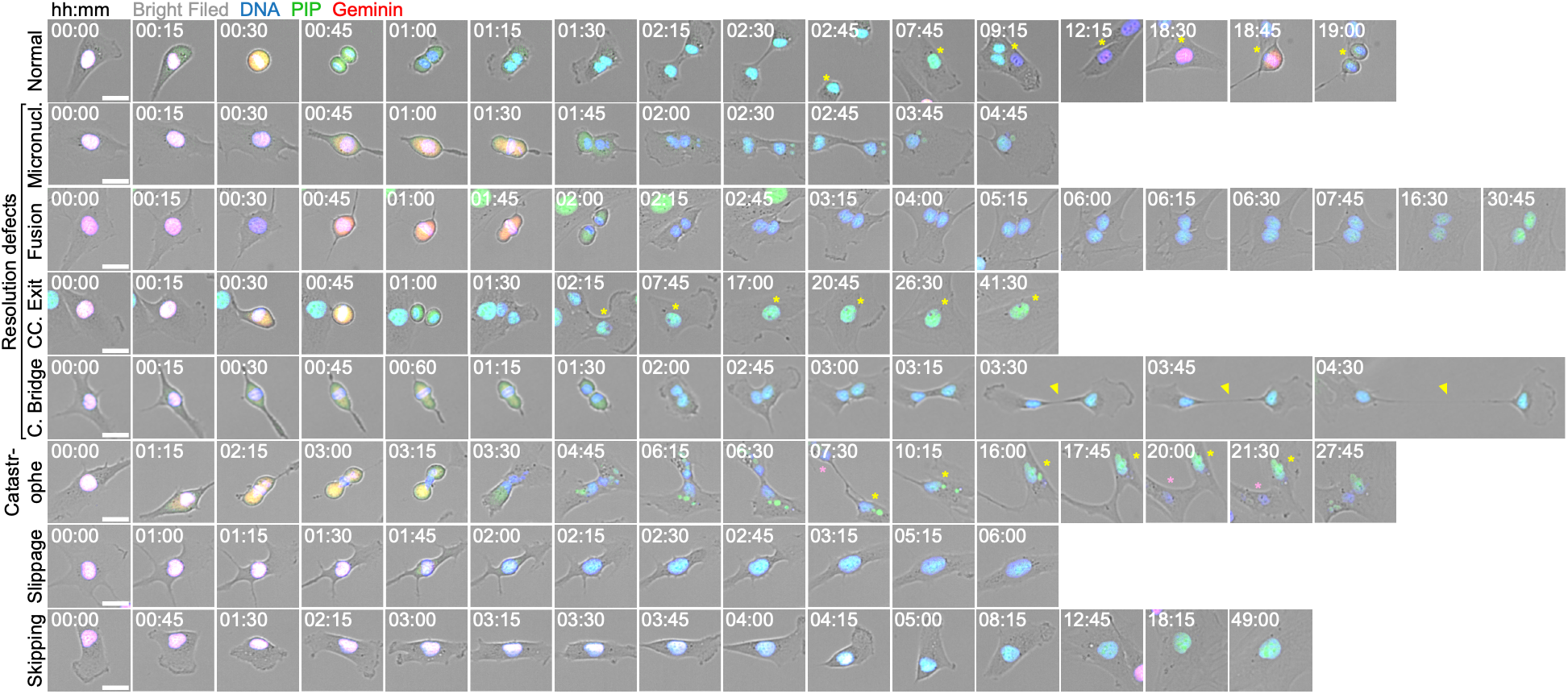
Documented mitotic defect classes in RPE-1 cells. Representative images illustrating mitotic defect categories detected in RPE-1 PIP-FUCCI cells: micronucleus formation, cellular fusion (failure of cytokinesis with fusion of daughter cells), cell-cycle exit (CC. Exit; post-mitotic arrest), cytoplasmic bridge (C. Bridge; persistent intercellular connection), mitotic catastrophe (abortive mitotic exit with apparent mechanical nuclear damage), mitotic slippage (exit from mitosis without anaphase), and mitotic skipping (G2-to-G1 transition without mitotic entry). Four-channel composite images show brightfield, DNA (SPY-DNA staining), PIP-eGFP and Geminin-mCherry. Examples were drawn from RPE-1 PIP-FUCCI cells under the perturbation conditions described in Figure 2. Scale bar, 25 µm.

**Supplementary Fig. S4.**
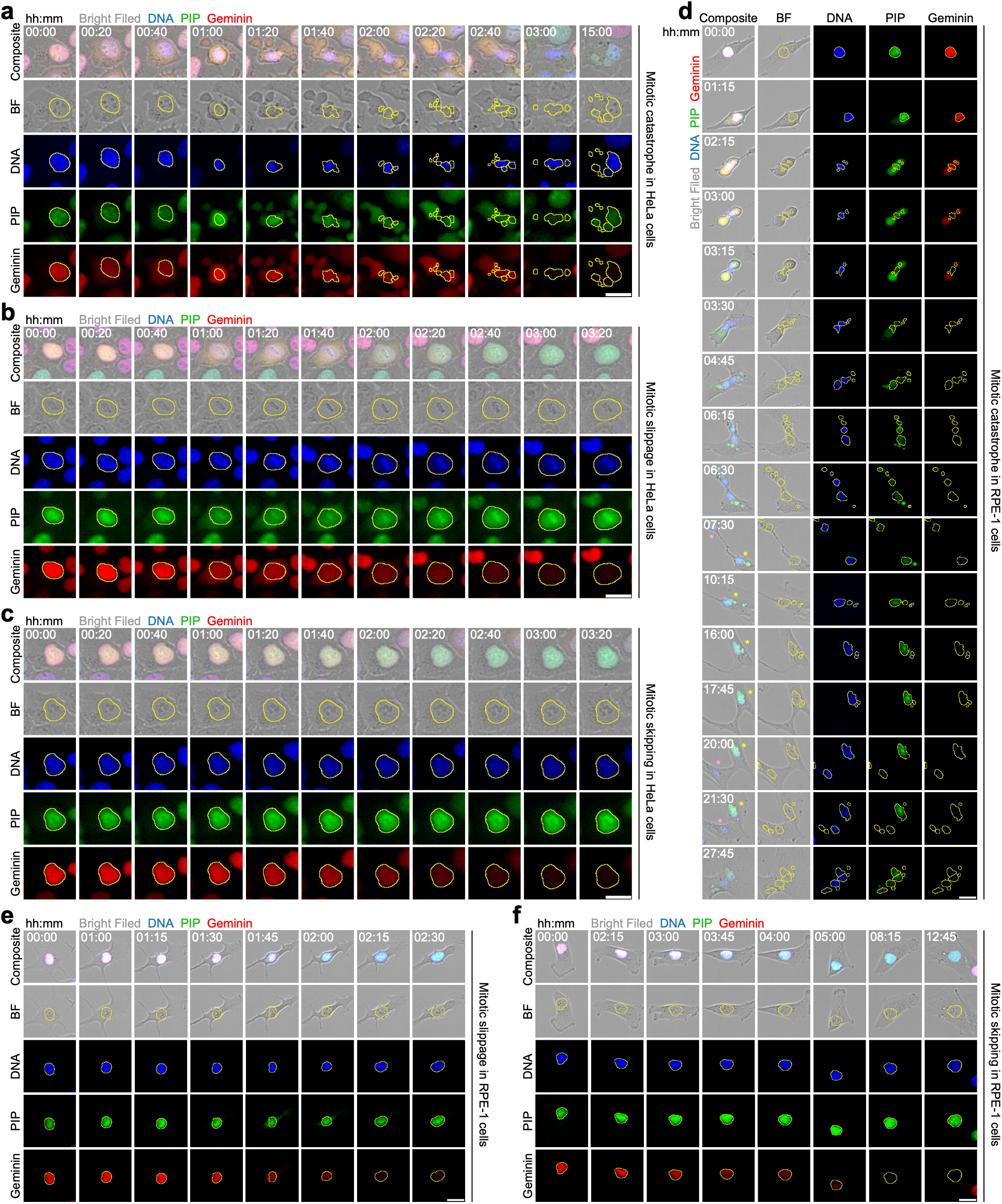
Mitotic catastrophe, slippage and skipping events in HeLa and RPE-1 cells. **(a-f)** Representative images of mitotic catastrophe, slippage and skipping events in HeLa (a-c) and RPE-1 (d-f) PIP-FUCCI cells. Images are also shown in Figure 2I and Supplementary Figure S2. Composites and individual single-channel views (brightfield, DNA, PIP-eGFP, Geminin-mCherry) are presented to clarify nuclear morphology. Yellow outlines indicate nuclear segmentation. Scale bar, 25 µm.

**Supplementary Fig. S5.**
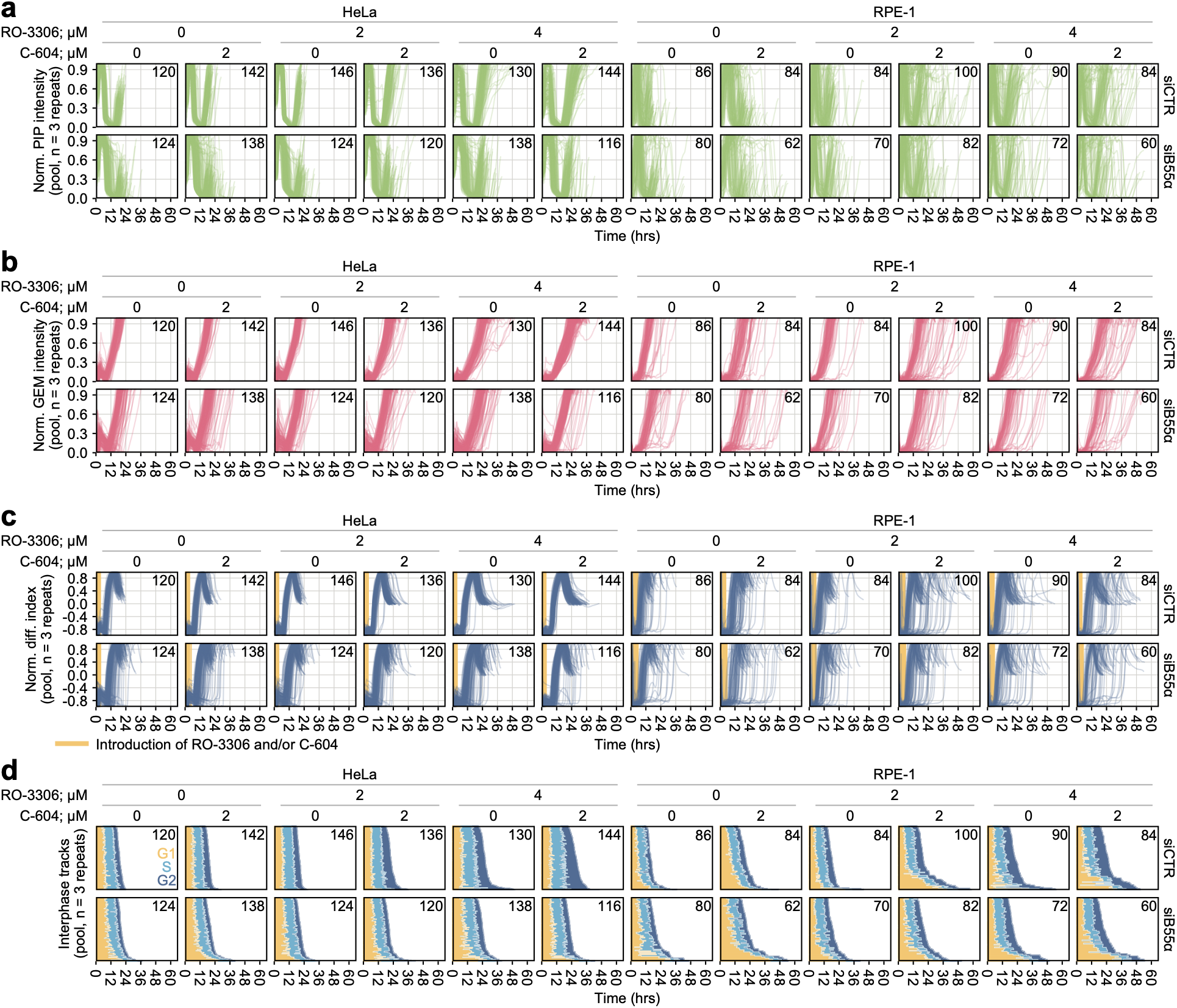
FUCCI analysis of interphase progression following perturbations of CDK1, GWL and PP2A-B55α. **(a, b)** Smoothed (moving average with a 9-bin window) min-max-normalised nuclear PIP-eGFP (a) and Geminin-mCherry (b) fluorescence intensities in tracked single HeLa and RPE-1 PIP-FUCCI cells under the indicated perturbation conditions. **(c)** Normalised difference index calculated as (Geminin − PIP) / (Geminin + PIP), where Geminin and PIP represent the smoothed normalised fluorescence intensities from (a, b). Yellow vertical lines mark the timing of the addition of RO-3306 or C-604. **(d)** Stratification of normalised difference indexes from (c) into G1, S and G2 phase segments per track, as described in (Fig. 2g, h). Tracks are ordered by total interphase duration. (a-d) Each line represents an individual interphase track. Results are pooled from n = 3 biological replicates. The total number of analysed tracks is shown in the upper-right corner of each panel.

**Supplementary Fig. S6.**
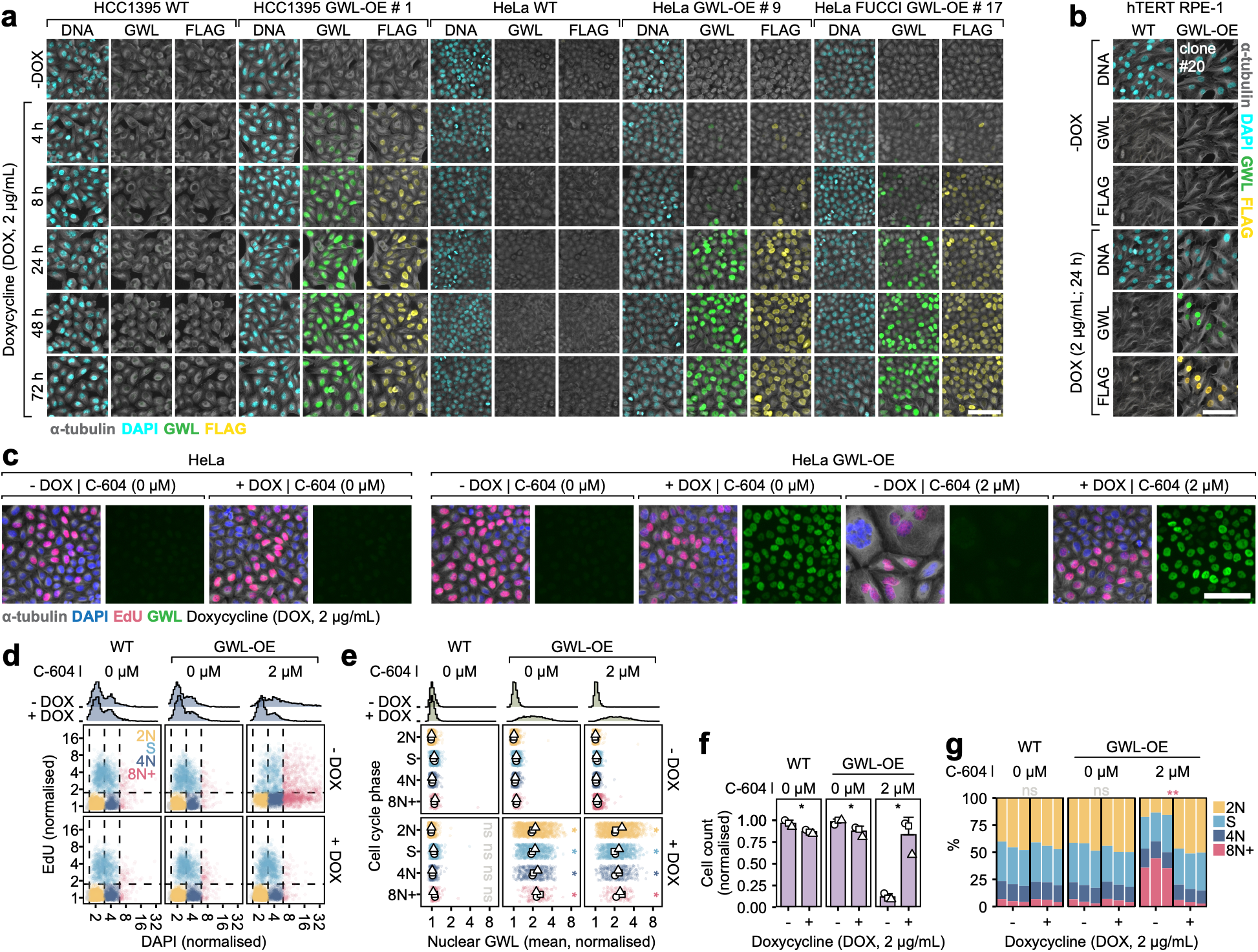
Development of an inducible GWL overexpression system. **(a, b)** Representative immunofluorescence images of HCC1395, HeLa (with and without PIP-FUCCI), and hTERT RPE-1 cell lines and their derived GWL-OE clones, grown without (−) or with (+) doxycycline (DOX; 2 µg/mL) for the indicated times. Scale bar is 50 µm. **(c)** Representative immunofluorescence images of HeLa GWL-OE cells under the four perturbation conditions (−/+DOX, 2 µg/mL; −/+C-604, 2 µM), alongside untreated parental HeLa cells (−/+DOX). DOX was added at time zero. C-604 was added 24 hours after DOX. Cells were fixed 72 hours after the addition of C-604. Scale bar, 100 µm. **(d)** Single-cell quantification of (c), shown as scatterplots of normalised integrated DAPI versus normalised mean nuclear EdU intensities. Cell-cycle phases 2N (G1), S (EdU-positive), 4N (G2/M), and polyploid (8N+) are colour-coded. Pooled results from n = 3 biological replicates (randomly subsampled 2,000 cells per condition per replicate). **(e)** Phase-specific quantification of GWL protein levels (normalised mean nuclear immunofluorescence intensity) following doxycycline treatment. Coloured points represent pooled single-cell measurements. White points and bars represent per-replicate medians (n = 3 biological replicates) and their means, respectively. Statistical significance was assessed using Welch’s two-tailed t-tests on per-replicate medians, comparing −DOX versus +DOX within each cell-cycle phase and C-604 treatment, with Benjamini-Hochberg correction across all 12 comparisons; ns – not significant, * p < 0.05. **(f)** Total cell counts normalised to the maximum count within each experiment. Bars and error bars represent means and standard deviations across n = 3 biological replicates, respectively, with individual values shown as points. Statistical significance was assessed using Welch’s two-tailed t-tests comparing −DOX versus +DOX within each cell line/C-604 condition, with Benjamini-Hochberg correction across the three comparisons. ns – not significant, * p < 0.05. **(g)** Proportions of 2N, S, 4N and polyploid 8N+ cells, classified as in (d), shown as three stacked bars representing results from n = 3 biological replicates. Statistical significance of differences in the 8N+ proportion was assessed using Welch’s two-tailed t-tests comparing −DOX versus +DOX within each cell line/C-604 condition, with Benjamini-Hochberg correction across the three comparisons. ns – not significant, ** p < 0.01.

**Supplementary Fig. S7.**
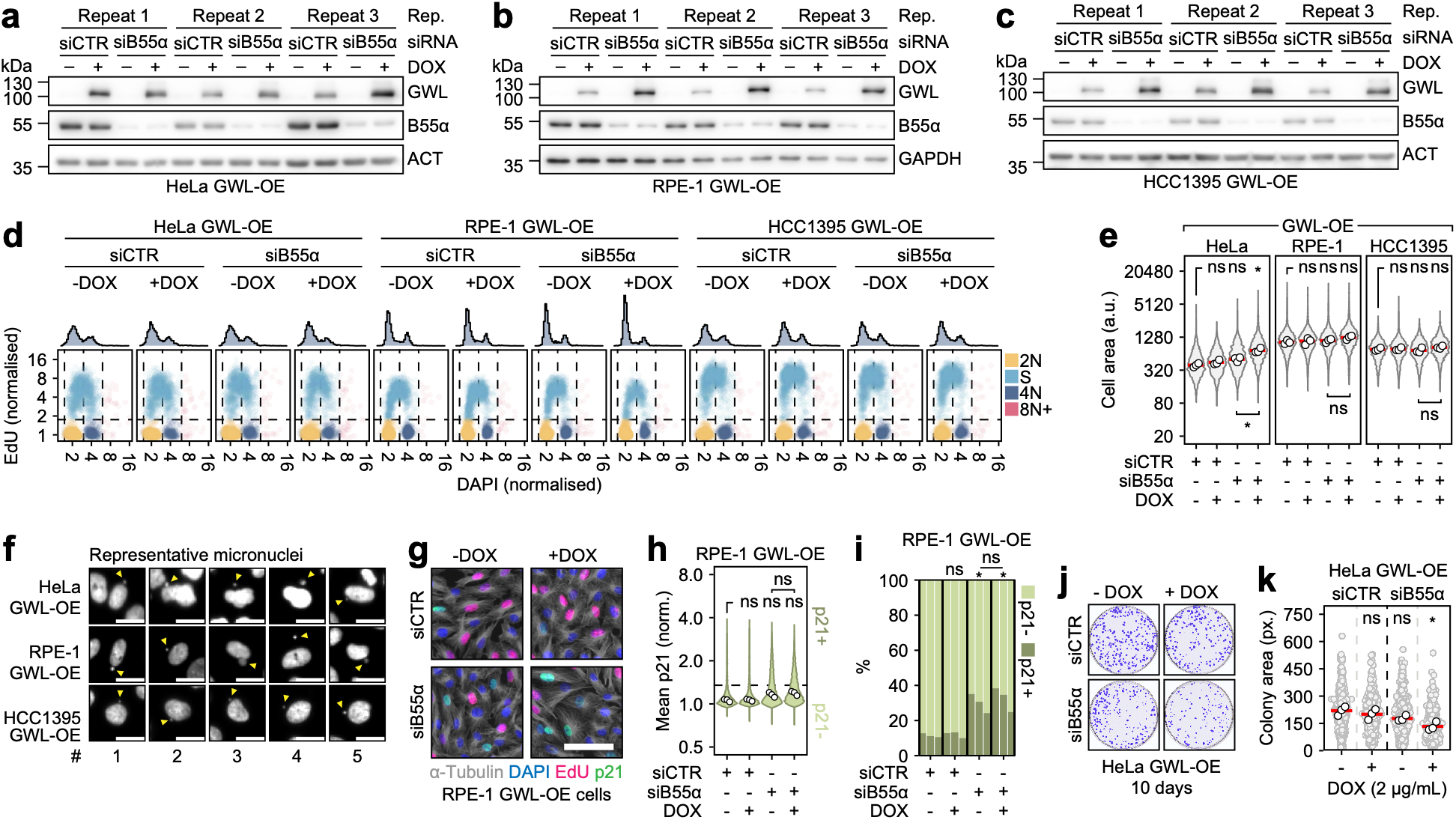
Extended characterisation of the combined effects of B55α depletion and GWL overexpression. **(a-c)** Western blots of GWL and B55α protein levels in HeLa (a), RPE-1 (b), and HCC1395 (c) GWL-OE cell lines, transfected with control (siCTR) or B55α-targeting (siB55α) siRNA and grown without (−) or with (+) doxycycline (DOX; 2 µg/mL, 24 hours, applied 24 hours post-transfection). Cells were harvested 48 hours post-transfection. β-actin (HeLa, HCC1395) or GAPDH (RPE-1) served as a loading control. Each blot shows n = 3 biological replicates. **(d)** Scatterplots of normalised integrated nuclear DAPI versus normalised mean nuclear EdU intensities in HeLa, RPE-1 and HCC1395 GWL-OE cells under four perturbation conditions (siCTR or siB55α, −/+DOX). Cells were transfected on day 0, treated with DOX (2 µg/mL) on day 1, and fixed on day 4. Cell-cycle phases (2N, S, 4N, 8N+) are colour-coded. Pooled data from n = 3 biological replicates (randomly subsampled 1,000 cells per condition per replicate). **(e)** Quantification of cell area in HeLa, RPE-1 and HCC1395 GWL-OE cells across the four perturbation conditions. Violin plots show distributions of pooled single-cell measurements. Points and horizontal red bars represent per-replicate medians (n = 3 biological replicates) and their means, respectively. Statistical significance was assessed using Welch’s two-tailed t-tests on per-replicate medians, comparing each treatment to the (siCTR, −DOX) control and (siB55α, −DOX) to (siB55α, +DOX), with p-values adjusted using the Benjamini-Hochberg procedure across the four comparisons per cell line. ns – not significant, * p < 0.05, ** p < 0.01. **(f)** Representative images of cells with micronuclei in HeLa, RPE-1 and HCC1395 GWL-OE cells under four perturbation conditions. Arrowheads indicate micronuclei. Scale bar represents 25 µm. **(g)** Representative composite immunofluorescence images (DAPI, EdU, α-tubulin, and p21) of RPE-1 GWL-OE cells across the four perturbation conditions. Scale bar represents 50 µm. **(h)** Normalised mean nuclear p21 intensity in RPE-1 GWL-OE cells, shown as violin plots of pooled single-cell measurements, with points representing per-replicate medians across n = 3 biological replicates. The threshold separating p21- and p21+ cells is indicated. **(i)** Proportions of p21- and p21+ cells, shown as three stacked bars representing results from n = 3 biological replicates. (h, i) Statistical significance was assessed using Welch’s two-tailed t-tests on medians (h) or proportions (i), comparing each treatment condition to the (siCTR, −DOX) control and (siB55α, −DOX) to (siB55α, +DOX), with Benjamini-Hochberg correction across the four comparisons per cell line. ns – not significant, * p < 0.05. **(J)** Representative images of colony formation assays in HeLa GWL-OE cells under the four perturbation conditions, stained with crystal violet. Cells were transfected, then seeded at low density 24 hours post-transfection in the presence (+) or absence (−) of DOX and cultured for 10 days before staining. **(K)** Colony size quantification. Grey points represent individual colony measurements pooled across n = 3 biological replicates. White points and red horizontal bars represent per-replicate medians and their means, respectively. Statistical significance was assessed using Welch’s two-tailed t-tests comparing the per-replicate medians of each condition with the control (siCTR, −DOX). The Benjamini-Hochberg correction was applied across the three comparisons. ns – not significant, * p < 0.05.

**Supplementary Fig. S8.**
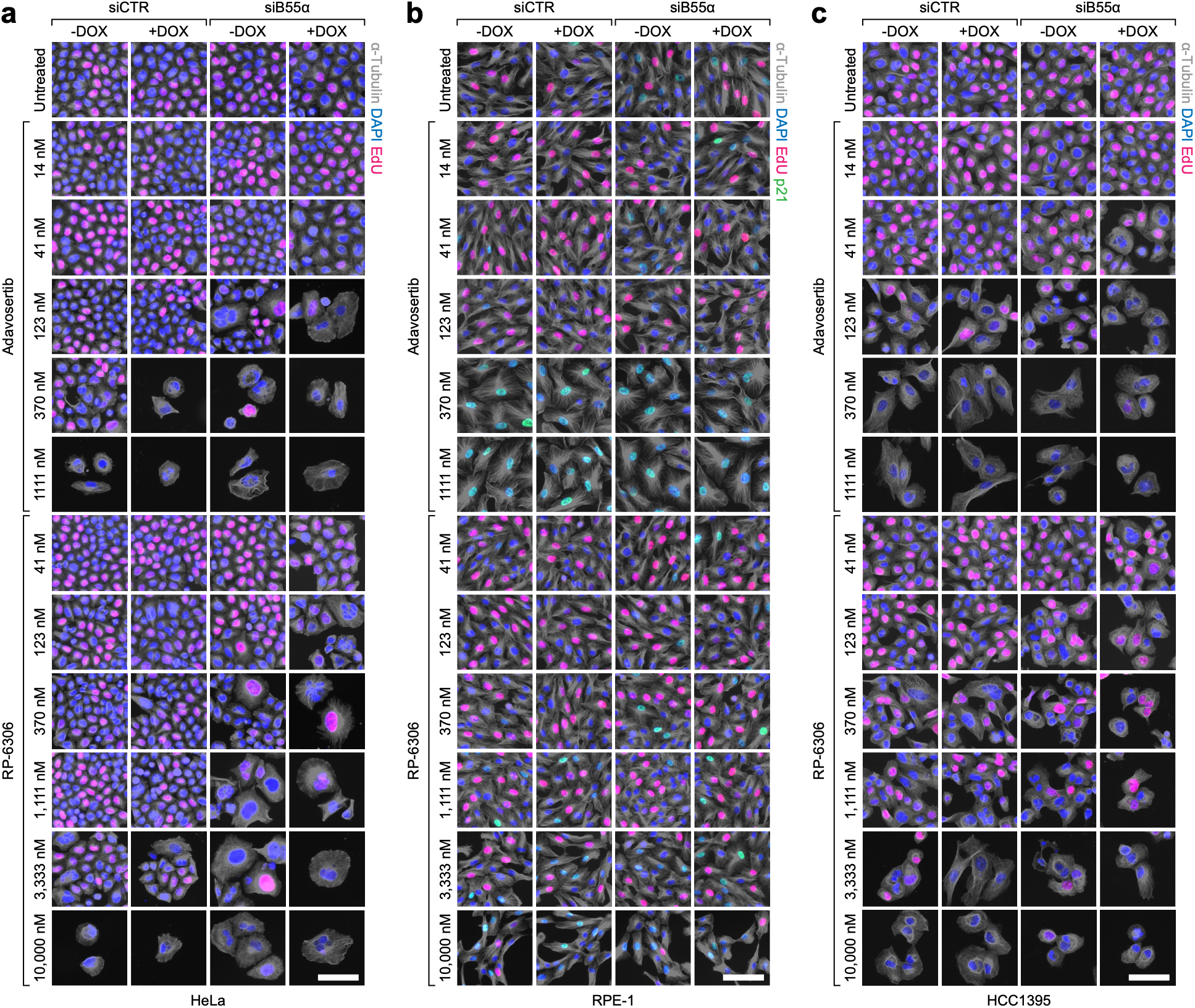
Dose-dependent effects of adavosertib and RP-6306 combined with B55α depletion and GWL overexpression on cellular physiology. **(a-c)** Representative composite immunofluorescence images of HeLa (a), RPE-1 (b) and HCC1395 (c) GWL-OE cells across the four perturbation conditions (siCTR or siB55α, −/+ DOX), treated with adavosertib (14 to 1,111 nM) or RP-6306 (41 to 10,000 nM). Composite images show DAPI, EdU, and α-tubulin in all three cell lines, with additional p21 staining in RPE-1. Scale bar, 25 µm.

**Supplementary Fig. S9.**
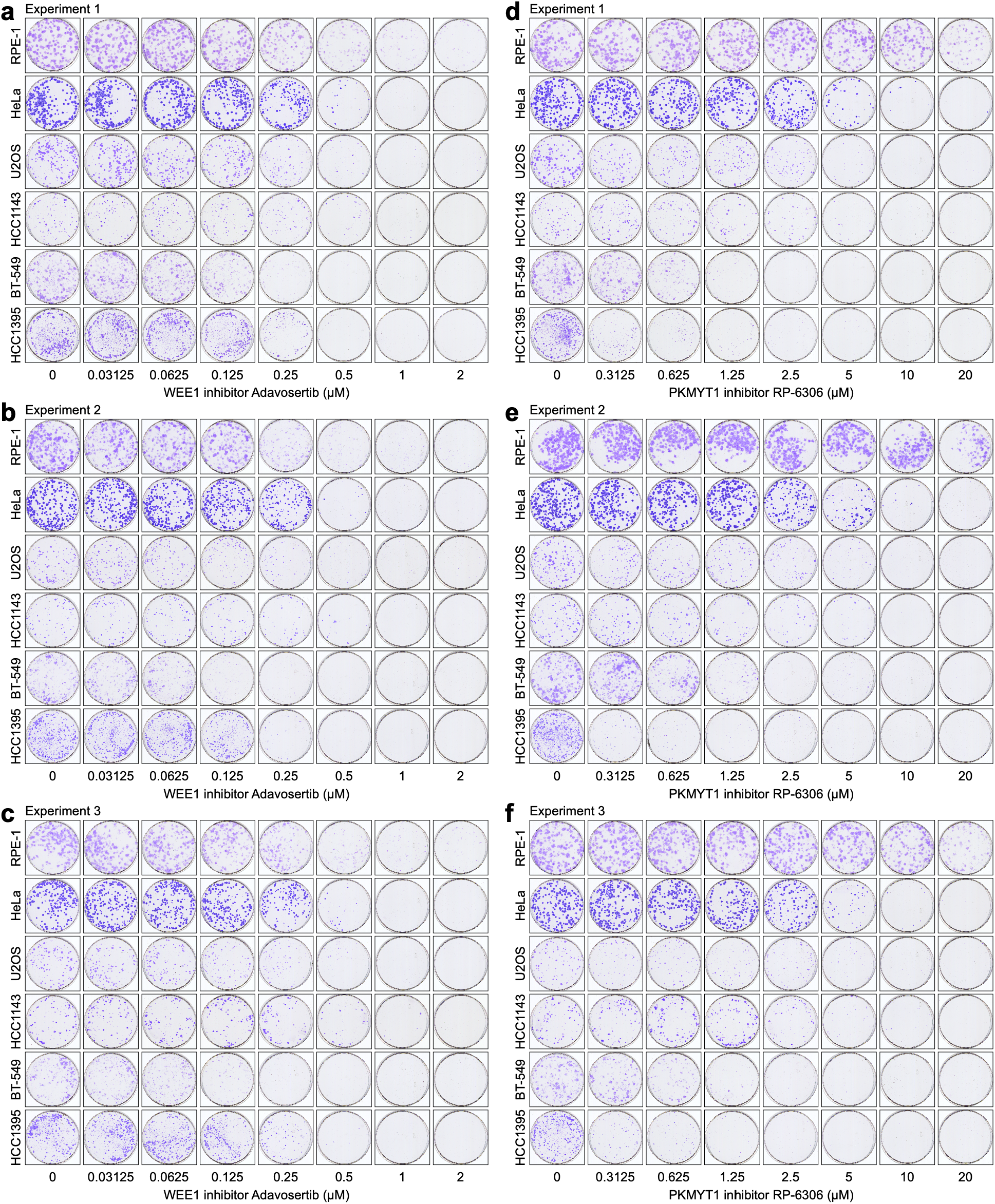
Dose-dependent effects of adavosertib and RP-6306 on clonogenic capacity. **(a-f)** Colonies (stained with crystal violet) of RPE-1, HeLa, U2OS, HCC1143, BT-549 and HCC1395 cells treated with increasing doses of adavosertib (a-c; 0 to 2 µM) and RP-6306 (d-f; 0 to 20 µM). Cells were grown in the presence of adavosertib or RP-6306 for 3 days, then cultured in drug-free medium for an additional 9-18 days, depending on the cell line.

**Supplementary Fig. S10.**
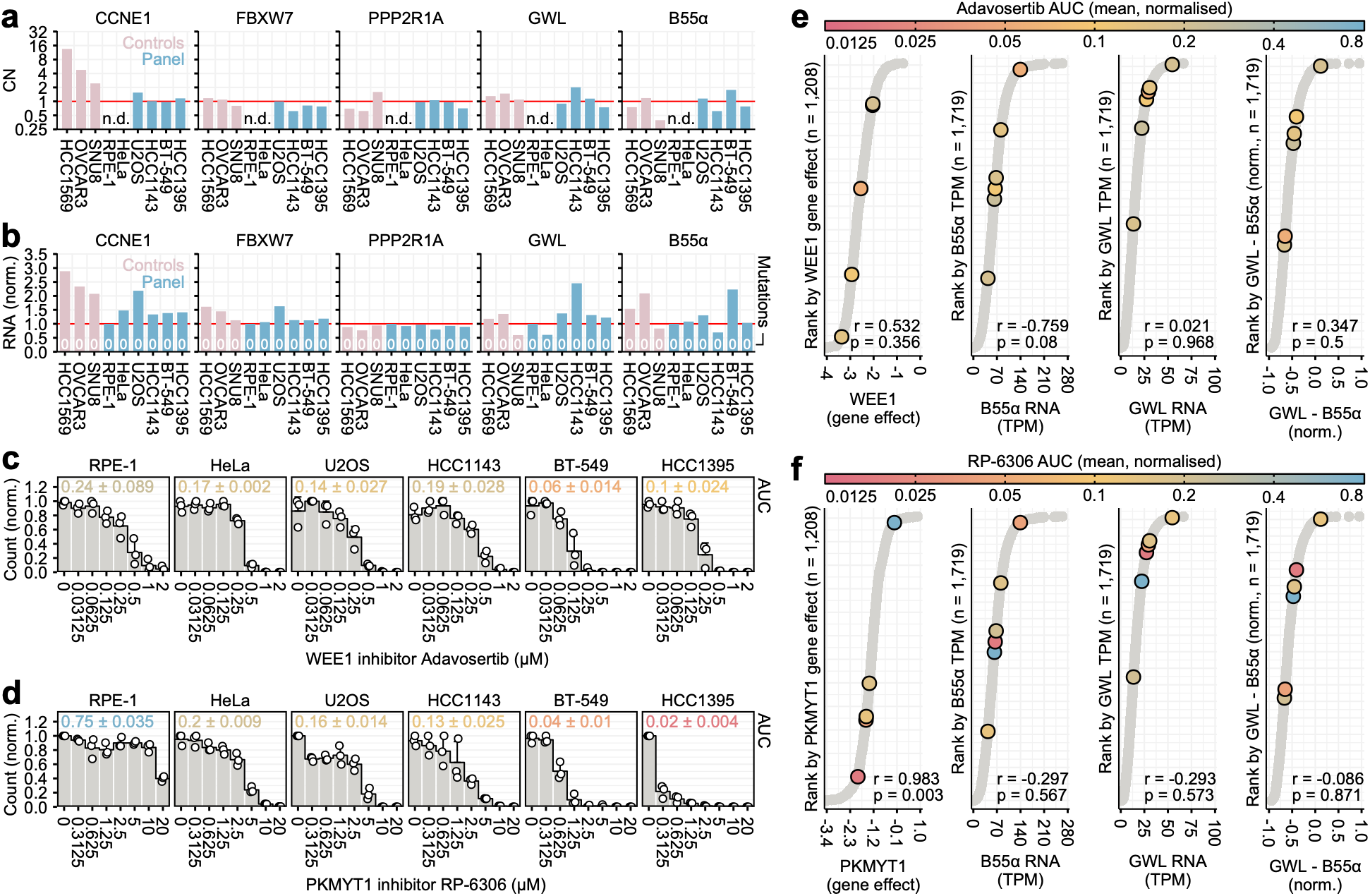
Assessment of B55α and GWL expression as biomarkers of cellular sensitivity to adavosertib and RP-6306. **(a-b)** DepMap (Cancer Dependency Map) copy number (a; linear scale, inferred from whole-genome sequencing) and RNA expression (b; TPM, normalised to RPE-1) of *CCNE1*, *FBXW7*, *PPP2R1A*, *MASTL* (*GWL*), and *PPP2R2A* (*B55α*). The number of damaging mutations in each gene is indicated in (b). Positive control cell lines (HCC1569, OVCAR3, SNU-8) carry known biomarkers of sensitivity to adavosertib and RP-6306 (*CCNE1* amplification, *FBXW7* loss). None of the experimental cell lines (RPE-1, HeLa, U2OS, HCC1143, BT-549, HCC1395) carries these biomarkers or damaging mutations in *PPP2R1A*, a proposed marker of sensitivity to PKMYT1 inhibition. **(c-d)** Quantification of colony formation capacity in the six experimental cell lines treated with adavosertib (c; 0-2 µM) and RP-6306 (d; 0-20 µM), expressed as colony counts normalised to the maximum within each cell line. Points represent results from n = 3 biological replicates. Bars and error bars represent means and standard deviations, respectively. Untreated controls produced between 71 ± 15 (HCC1143) and 223 ± 23 (HeLa) colonies. AUC values, normalised so that AUC = 1 indicates no drug effect across the dose range, are shown for each cell line. **(e-f)** Relationship between Adavosertib (e) and RP-6306 (f) sensitivity and four candidate biomarker rankings. Each subpanel ranks DepMap cell lines (grey points; n indicated per subpanel) by a candidate biomarker: WEE1 (e) or PKMYT1 (f) gene effect (CRISPR Chronos dependency score), B55α TPM, GWL TPM, and the normalised GWL-B55α difference. The PKMYT1 subpanel reflects the subset of DepMap cell lines with available CRISPR dependency data (n = 1,208), whereas the other subpanels include all cell lines with RNA expression data (n = 1,719). The experimental cell lines from (c, d) are overlaid as larger coloured points, with colours indicating their mean normalised AUC values for Adavosertib (e) and RP-6306 (f). Pearson correlation coefficients (r) and p-values quantify the relationship between each candidate biomarker ranking and the experimental AUC values.

**Supplementary Fig. S11.**
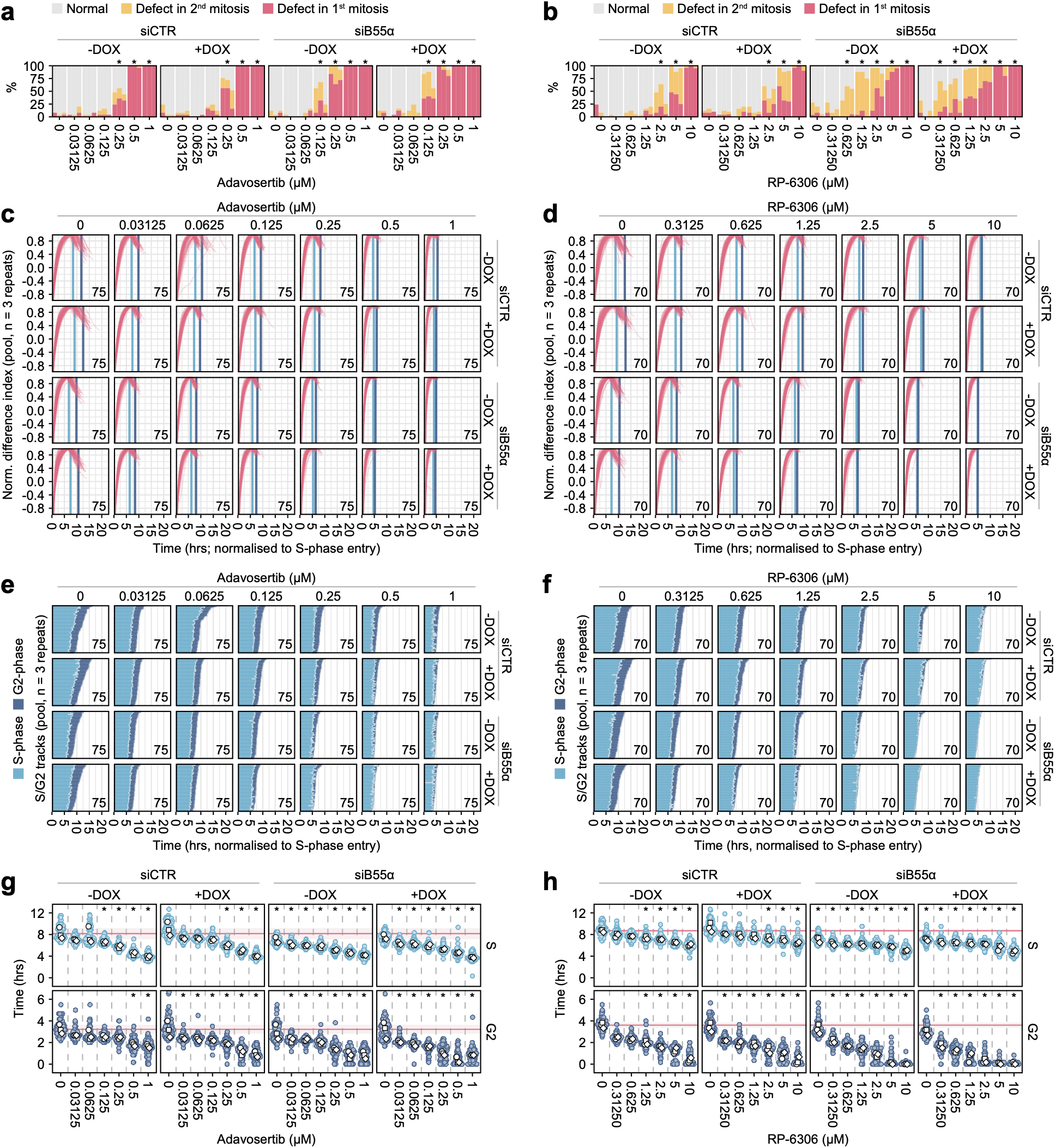
PIP-FUCCI analysis of cell cycle progression following perturbations of GWL and PP2A-B55α, combined with WEE1 and PKMYT1 inhibition. **(a, b)** Proportions of normal and defective (1st or 2nd cell division) mitoses in HeLa PIP-FUCCI GWL-OE cells across four perturbation conditions (siCTR or siB55α, −/+DOX), treated with adavosertib (a; 0 to 1 µM) or RP-6306 (b; 0 to 10 µM). For each condition, proportions are shown as three stacked bars, representing results from n = 3 biological replicates. Statistical significance was assessed using a linear model with treatment condition as a categorical predictor, followed by Dunnett’s post hoc test comparing each treatment to the (siCTR, −DOX, untreated) control. Significance is shown qualitatively. * p < 0.05. **(c, d)** Dose-dependent effects of adavosertib (c) and RP-6306 (d) on normalised difference indices, calculated as (Geminin − PIP) / (Geminin + PIP), where Geminin and PIP denote smoothed normalised fluorescence intensities. Pooled results from n = 3 biological replicates are shown, with 20-25 tracks per condition per replicate. The total number of tracks is indicated in the bottom-right corners. Only S and G2 phases are shown, with t = 0 representing S-phase entry. Vertical light-blue and dark-blue lines mark the median S/G2 and G2/M transitions, respectively. **(e, f)** Stratification of normalised difference indices from (c, d) into S and G2 phase segments per track for adavosertib (e) and RP-6306 (f), as in (Fig. 2g, h). **(g, h)** Quantification of S and G2 durations following acute adavosertib (g) or RP-6306 (h) treatment, derived from (E, F). Coloured and white points represent pooled single-cell measurements from n = 3 biological replicates and their per-replicate medians, respectively. Horizontal red lines and shaded regions represent the overall means and standard deviations of the untreated control (siCTR, −DOX, untreated). Statistical significance was assessed using the Kruskal-Wallis test, followed by Dunn’s post hoc test with Bonferroni correction for all pairwise comparisons; only comparisons with the untreated control are displayed. Significance is shown qualitatively. * p < 0.01.

## Descriptions of Additional Supplementary Movies

**Supplementary Movie M1.**

**PIP-FUCCI interphase analysis.** Automated stratification of 100 representative PIP-FUCCI interphase tracks randomly selected from a pool of n = 2,532 tracks.

**Supplementary Movie M2.**

**Graded mitotic defects in HeLa cells.** HeLa PIP-FUCCI cells grown under normal conditions (0-16 h) and then treated with different doses of the GWL inhibitor C-604 and the CDK1 inhibitor RO-3306 (C-604 | RO-3306; 16-64 h). Composite images of four channels (Grey – bright field, Red – Geminin, Green – PIP, Blue – DNA) are shown. The scale bar is 100 µm.

**Supplementary Movie M3.**

**Graded mitotic defects in hTERT RPE-1 cells.** RPE-1 PIP-FUCCI cells grown under normal conditions (0-6 h) and then treated with different doses of the GWL inhibitor C-604 and the CDK1 inhibitor RO-3306 (C-604 | RO-3306; 6-72 h). Composite images of four channels (Grey – bright field, Red – Geminin, Green – PIP, Blue – DNA) are shown. The scale bar is 100 µm.

**Supplementary Movie M4.**

**The impact of WEE1 inhibition on HeLa cells with low B55α and/or high GWL levels.** HeLa PIP-FUCCI GWL-OE cells transfected with siCTR or siB55α, grown without (−) or with (+) doxycycline (DOX), and treated with the indicated concentrations (0-1 µM) of the WEE1 inhibitor adavosertib. Composite images of four channels (Grey – bright field, Red – Geminin, Green – PIP, Blue – DNA) are shown. The scale bar is 50 µm.

**Supplementary Movie M5.**

**The impact of PKMYT1 inhibition on HeLa cells with low B55α and/or high GWL levels.** HeLa PIP-FUCCI GWL-OE cells transfected with siCTR or siB55α, grown without (−) or with (+) doxycycline (DOX), and treated with the indicated concentrations (0-1 µM) of the PKMYT1 inhibitor RP-6306. Composite images of four channels (Grey – bright field, Red – Geminin, Green – PIP, Blue – DNA) are shown. The scale bar is 50 µm.

## References

1. Novák, B. & Tyson, J. J. Mechanisms of signalling-memory governing progression through the eukaryotic cell cycle. Curr. Opin. Cell Biol. 69, 7–16 (2021).

2. Tyson, J. J. & Novak, B. Temporal organization of the cell cycle. Curr. Biol. 18, R759– R768 (2008).

3. Crncec, A. & Hochegger, H. Triggering mitosis. FEBS Lett. 593, 2868–2888 (2019).

4. Gavet, O. & Pines, J. Progressive activation of CyclinB1-Cdk1 coordinates entry to mitosis. Dev. Cell 18, 533–43 (2010).

5. Parker, L. L. & Piwnica-Worms, H. Inactivation of the p34cdc2-cyclin B complex by the human WEE1 tyrosine kinase. Science 257, 1955–7 (1992).

6. Russell, P. & Nurse, P. Negative regulation of mitosis by wee1+, a gene encoding a protein kinase homolog. Cell 49, 559–67 (1987).

7. Liu, F., Stanton, J. J., Wu, Z. & Piwnica-Worms, H. The human Myt1 kinase preferentially phosphorylates Cdc2 on threonine 14 and localizes to the endoplasmic reticulum and Golgi complex. Mol. Cell. Biol. 17, 571–83 (1997).

8. Booher, R. N., Holman, P. S. & Fattaey, A. Human Myt1 is a cell cycle-regulated kinase that inhibits Cdc2 but not Cdk2 activity. J. Biol. Chem. 272, 22300–6 (1997).

9. Schmitz, M. H. A. et al. Live-cell imaging RNAi screen identifies PP2A-B55alpha and importin-beta1 as key mitotic exit regulators in human cells. Nat. Cell Biol. 12, 886–93 (2010).

10. Cundell, M. J. et al. A PP2A-B55 recognition signal controls substrate dephosphorylation kinetics during mitotic exit. J. Cell Biol. 214, 539–54 (2016).

11. Kruse, T. et al. Substrate recognition principles for the PP2A-B55 protein phosphatase. Sci. Adv. 10, eadp5491 (2024).

12. Kumagai, A. & Dunphy, W. G. The cdc25 protein controls tyrosine dephosphorylation of the cdc2 protein in a cell-free system. Cell 64, 903–14 (1991).

13. Gautier, J., Solomon, M. J., Booher, R. N., Bazan, J. F. & Kirschner, M. W. cdc25 is a specific tyrosine phosphatase that directly activates p34cdc2. Cell 67, 197–211 (1991).

14. Coleman, T. R. & Dunphy, W. G. Cdc2 regulatory factors. Curr. Opin. Cell Biol. 6, 877–82 (1994).

15. Perry, J. A. & Kornbluth, S. Cdc25 and Wee1: analogous opposites? Cell Div. 2, 12 (2007).

16. Watanabe, N., et al. Cyclin-Dependent Kinase (CDK) Phosphorylation Destabilizes Somatic Wee1 via Multiple Pathways. www.pnas.orgcgidoi10.1073pnas.0500410102 (2005).

17. Blake-Hodek, K. A. et al. Determinants for activation of the atypical AGC kinase Greatwall during M phase entry. Mol. Cell. Biol. 32, 1337–53 (2012).

18. Vigneron, S. et al. Characterization of the mechanisms controlling Greatwall activity. Mol. Cell. Biol. 31, 2262–75 (2011).

19. Mochida, S., Maslen, S. L., Skehel, M. & Hunt, T. Greatwall phosphorylates an inhibitor of protein phosphatase 2A that is essential for mitosis. Science 330, 1670–3 (2010).

20. Gharbi-Ayachi, A. et al. The substrate of Greatwall kinase, Arpp19, controls mitosis by inhibiting protein phosphatase 2A. Science 330, 1673–7 (2010).

21. Williams, B. C. et al. Greatwall-phosphorylated Endosulfine is both an inhibitor and a substrate of PP2A-B55 heterotrimers. Elife 3, e01695 (2014).

22. Padi, S. K. R. et al. Cryo-EM structures of PP2A:B55-FAM122A and PP2A:B55-ARPP19. Nature 625, 195–203 (2024).

23. Dephoure, N. et al. A quantitative atlas of mitotic phosphorylation. Proc. Natl. Acad. Sci. U. S. A. 105, 10762–7 (2008).

24. Olsen, J. V et al. Quantitative phosphoproteomics reveals widespread full phosphorylation site occupancy during mitosis. Sci. Signal. 3, ra3 (2010).

25. Linder, M. I. et al. Mitotic Disassembly of Nuclear Pore Complexes Involves CDK1- and PLK1-Mediated Phosphorylation of Key Interconnecting Nucleoporins. Dev. Cell 43, 141–156.e7 (2017).

26. Zach, R. et al. The balance between B55α and Greatwall expression levels predicts sensitivity to Greatwall inhibition in cancer cells. Nat. Commun. 16, 8016 (2025).

27. Peters, J.-M. The anaphase promoting complex/cyclosome: a machine designed to destroy. Nat. Rev. Mol. Cell Biol. 7, 644–56 (2006).

28. Holder, J., Mohammed, S. & Barr, F. A. Ordered dephosphorylation initiated by the selective proteolysis of cyclin B drives mitotic exit. Elife 9, 1–33 (2020).

29. Hégarat, N. et al. PP2A/B55 and Fcp1 regulate Greatwall and Ensa dephosphorylation during mitotic exit. PLoS Genet. 10, e1004004 (2014).

30. Cundell, M. J. et al. The BEG (PP2A-B55/ENSA/Greatwall) pathway ensures cytokinesis follows chromosome separation. Mol. Cell 52, 393–405 (2013).

31. McCloy, R. A. et al. Global Phosphoproteomic Mapping of Early Mitotic Exit in Human Cells Identifies Novel Substrate Dephosphorylation Motifs. Mol. Cell. Proteomics 14, 2194–212 (2015).

32. Vassilev, L. T. et al. Selective small-molecule inhibitor reveals critical mitotic functions of human CDK1. Proc. Natl. Acad. Sci. U. S. A. 103, 10660–5 (2006).

33. McCloy, R. A. et al. Partial inhibition of Cdk1 in G2 phase overrides the SAC and decouples mitotic events. Cell Cycle 13, 1400–1412 (2014).

34. Hochegger, H. et al. An essential role for Cdk1 in S phase control is revealed via chemical genetics in vertebrate cells. J. Cell Biol. 178, 257–68 (2007).

35. Ma, H. T., Tsang, Y. H., Marxer, M. & Poon, R. Y. C. Cyclin A2-cyclin-dependent kinase 2 cooperates with the PLK1-SCFbeta-TrCP1-EMI1-anaphase-promoting complex/cyclosome axis to promote genome reduplication in the absence of mitosis. Mol. Cell. Biol. 29, 6500–14 (2009).

36. Gravells, P., Tomita, K., Booth, A., Poznansky, J. & Porter, A. C. G. Chemical genetic analyses of quantitative changes in Cdk1 activity during the human cell cycle. Hum. Mol. Genet. 22, 2842–51 (2013).

37. Vassilev, L. T. Cell cycle synchronization at the G2/M phase border by reversible inhibition of CDK1. Cell Cycle 5, 2555–6 (2006).

38. Voets, E., Marsman, J., Demmers, J., Beijersbergen, R. & Wolthuis, R. The lethal response to Cdk1 inhibition depends on sister chromatid alignment errors generated by KIF4 and isoform 1 of PRC1. Sci. Rep. 5, (2015).

39. Yu, J. et al. Greatwall kinase: a nuclear protein required for proper chromosome condensation and mitotic progression in Drosophila. J. Cell Biol. 164, 487–92 (2004).

40. Voets, E. & Wolthuis, R. M. F. MASTL is the human orthologue of Greatwall kinase that facilitates mitotic entry, anaphase and cytokinesis. Cell Cycle 9, 3591–601 (2010).

41. Burgess, A. et al. Loss of human Greatwall results in G2 arrest and multiple mitotic defects due to deregulation of the cyclin B-Cdc2/PP2A balance. Proc. Natl. Acad. Sci. U. S. A. 107, 12564–9 (2010).

42. Álvarez-Fernández, M. et al. Greatwall is essential to prevent mitotic collapse after nuclear envelope breakdown in mammals. Proc. Natl. Acad. Sci. U. S. A. 110, 17374– 9 (2013).

43. Sanz-Flores, M. et al. PP2A-B55 phosphatase counteracts Ki-67-dependent chromosome individualization during mitosis. Cell Rep. 43, 114494 (2024).

44. Amin, P. et al. PP2A-B55: substrates and regulators in the control of cellular functions. Oncogene 41, 1–14 (2022).

45. Mailand, N. et al. Regulation of G(2)/M events by Cdc25A through phosphorylation-dependent modulation of its stability. EMBO J. 21, 5911–20 (2002).

46. Boutros, R., Lobjois, V. & Ducommun, B. CDC25 phosphatases in cancer cells: key players? Good targets? Nat. Rev. Cancer 7, 495–507 (2007).

47. Wang, Q., Bode, A. M. & Zhang, T. Targeting CDK1 in cancer: mechanisms and implications. *NPJ Precis*. Oncol. 7, 58 (2023).

48. Suski, J. M., Braun, M., Strmiska, V. & Sicinski, P. Targeting cell-cycle machinery in cancer. Cancer Cell 39, 759–778 (2021).

49. Rogers, S. et al. MASTL overexpression promotes chromosome instability and metastasis in breast cancer. Oncogene 37, 4518–4533 (2018).

50. Vera, J. et al. Greatwall promotes cell transformation by hyperactivating AKT in human malignancies. Elife 4, (2015).

51. Fatima, I. et al. MASTL regulates EGFR signaling to impact pancreatic cancer progression. Oncogene 40, 5691–5704 (2021).

52. Choi, K.-M. et al. Activity-based protein profiling and global proteome analysis reveal MASTL as a potential therapeutic target in gastric cancer. Cell Commun. Signal. 22, 397 (2024).

53. Goguet-Rubio, P. et al. PP2A-B55 Holoenzyme Regulation and Cancer. Biomolecules 10, 1–17 (2020).

54. Curtis, C. et al. The genomic and transcriptomic architecture of 2,000 breast tumours reveals novel subgroups. Nature 486, 346–52 (2012).

55. Cheng, Y. et al. Evaluation of PPP2R2A as a prostate cancer susceptibility gene: a comprehensive germline and somatic study. Cancer Genet. 204, 375–81 (2011).

56. Beca, F., Pereira, M., Cameselle-Teijeiro, J. F., Martins, D. & Schmitt, F. Altered PPP2R2A and Cyclin D1 expression defines a subgroup of aggressive luminal-like breast cancer. BMC Cancer 15, 285 (2015).

57. Zhao, Z. et al. PPP2R2A prostate cancer haploinsufficiency is associated with worse prognosis and a high vulnerability to B55α/PP2A reconstitution that triggers centrosome destabilization. Oncogenesis 8, 72 (2019).

58. Qiu, Z. et al. PPP2R2A insufficiency enhances PD-L1 immune checkpoint blockade efficacy in lung cancer through cGAS-STING activation. J. Clin. Invest. 136, (2026).

59. Kalev, P. et al. Loss of PPP2R2A inhibits homologous recombination DNA repair and predicts tumor sensitivity to PARP inhibition. Cancer Res. 72, 6414–24 (2012).

60. Grant, G. D., Kedziora, K. M., Limas, J. C., Cook, J. G. & Purvis, J. E. Accurate delineation of cell cycle phase transitions in living cells with PIP-FUCCI. Cell Cycle 17, 2496–2516 (2018).

61. Sakaue-Sawano, A. et al. Visualizing spatiotemporal dynamics of multicellular cell-cycle progression. Cell 132, 487–98 (2008).

62. Liao, H. et al. CDK1 promotes nascent DNA synthesis and induces resistance of cancer cells to DNA-damaging therapeutic agents. Oncotarget 8, 90662–90673 (2017).

63. Lindqvist, A., van Zon, W., Karlsson Rosenthal, C. & Wolthuis, R. M. F. Cyclin B1-Cdk1 activation continues after centrosome separation to control mitotic progression. PLoS Biol. 5, e123 (2007).

64. Álvarez-Fernández, M. et al. Therapeutic relevance of the PP2A-B55 inhibitory kinase MASTL/Greatwall in breast cancer. Cell Death Differ. 25, 828–840 (2018).

65. Hirai, H. et al. Small-molecule inhibition of Wee1 kinase by MK-1775 selectively sensitizes p53-deficient tumor cells to DNA-damaging agents. Mol. Cancer Ther. 8, 2992–3000 (2009).

66. Szychowski, J. et al. Discovery of an Orally Bioavailable and Selective PKMYT1 Inhibitor, RP-6306. J. Med. Chem. 65, 10251–10284 (2022).

67. Wong, P. Y., Ma, H. T., Lee, H. & Poon, R. Y. C. MASTL(Greatwall) regulates DNA damage responses by coordinating mitotic entry after checkpoint recovery and APC/C activation. Sci. Rep. 6, 22230 (2016).

68. Gobran, M. et al. PLK1 inhibition delays mitotic entry revealing changes to the phosphoproteome of mammalian cells early in division. EMBO J. 44, 1891–1920 (2025).

69. Kettenbach, A. N. et al. Quantitative phosphoproteomics identifies substrates and functional modules of Aurora and Polo-like kinase activities in mitotic cells. Sci. Signal. 4, rs5 (2011).

70. Hartwell, L. H. & Weinert, T. A. Checkpoints: controls that ensure the order of cell cycle events. Science 246, 629–34 (1989).

71. Rhind, N. & Russell, P. Signaling pathways that regulate cell division. Cold Spring Harb. Perspect. Biol. 4, (2012).

72. Belbelazi, A. et al. PKMYT1 has an important role in the timing and fidelity of chromosome segregation. EMBO Rep. 27, 3564–3584 (2026).

73. Willaume, S. et al. CRISPR-PTM and CRISPR-VEIS: Multiplexed platforms for quantitative functional analysis of endogenous phosphosites. Preprint at 10.64898/2026.05.07.723463 (2026).

74. Elbæk, C. R. et al. WEE1 kinase protects the stability of stalled DNA replication forks by limiting CDK2 activity. Cell Rep. 38, 110261 (2022).

75. Beck, H. et al. Cyclin-dependent kinase suppression by WEE1 kinase protects the genome through control of replication initiation and nucleotide consumption. Mol. Cell. Biol. 32, 4226–36 (2012).

76. Domínguez-Kelly, R. et al. Wee1 controls genomic stability during replication by regulating the Mus81-Eme1 endonuclease. J. Cell Biol. 194, 567–79 (2011).

77. Duda, H. et al. A Mechanism for Controlled Breakage of Under-replicated Chromosomes during Mitosis. Dev. Cell 39, 740–755 (2016).

78. Yap, T. A. et al. Abstract PR008: MYTHIC: First-in-human (FIH) biomarker-driven phase I trial of PKMYT1 inhibitor lunresertib (lunre) alone and with ATR inhibitor camonsertib (cam) in solid tumors with *CCNE1* amplification or deleterious alterations in *FBXW7* or *PPP2R1A*. Mol. Cancer Ther. 22, PR008–PR008 (2023).

79. Gallo, D. et al. CCNE1 amplification is synthetic lethal with PKMYT1 kinase inhibition. Nature 604, 749–756 (2022).

80. Schram, A. M. et al. Abstract CT262: Efficacy and safety of the combination PKMYT1-inhibitor lunresertib and ATR-inhibitor camonsertib in patients with ovarian and endometrial cancers: Phase I MYTHIC study (NCT04855656). Cancer Res. 85, CT262–CT262 (2025).

81. Mátés, L. et al. Molecular evolution of a novel hyperactive Sleeping Beauty transposase enables robust stable gene transfer in vertebrates. Nat. Genet. 41, 753– 61 (2009).

82. Schneider, C. A., Rasband, W. S. & Eliceiri, K. W. NIH Image to ImageJ: 25 years of image analysis. Nat. Methods 9, 671–5 (2012).

83. Allan, C. et al. OMERO: flexible, model-driven data management for experimental biology. Nat. Methods 9, 245–53 (2012).

84. Stringer, C., Wang, T., Michaelos, M. & Pachitariu, M. Cellpose: a generalist algorithm for cellular segmentation. Nat. Methods 18, 100–106 (2021).

85. Gallusser, B. & Weigert, M. Trackastra: Transformer-based cell tracking for live-cell microscopy. http://arxiv.org/abs/2405.15700 (2024).

